# Single-cell transcriptomic landscape of the neuroimmune compartment in amyotrophic lateral sclerosis brain and spinal cord

**DOI:** 10.1101/2024.11.12.623183

**Authors:** John F. Tuddenham, Masashi Fujita, Anthony Khairallah, Claire Harbison, Xena E. Flowers, Guillermo Coronas-Samano, Silas Maniatis, Aidan Daly, Julie A. Schneider, Andrew F. Teich, Jean Paul G. Vonsattel, Peter A. Sims, Wassim Elyaman, Elizabeth M. Bradshaw, Hemali Phatnani, Neil Shneider, David A. Bennett, Philip L. De Jager, Serge Przedborski, Vilas Menon, Marta Olah

**Affiliations:** Department of Neurology, Columbia University Medical Center, New York, NY, USA; Center for Translational & Computational Neuroimmunology, Department of Neurology, Columbia University Irving Medical Center, New York, NY, USA; Taub Institute for Research on Alzheimer’s Disease and the Aging Brain, Columbia University Medical Center, New York, NY, USA, Department of Pathology and Cell Biology, Columbia University Medical Center, New York, NY, USA; Rush Alzheimer’s Disease Center, Rush University Medical Center, Chicago, IL, USA; Department of Systems Biology, Columbia University Medical Center, New York, NY, USA; Department of Biochemistry and Molecular Biophysics, Columbia University Medical Center, New York, NY, USA; Eleanor and Lou Gehrig ALS Center, Columbia University Medical Center, New York, USA; Center for Motor Neuron Biology and Disease, Columbia University Medical Center, New York, NY, USA; Center for Genomics of Neurodegenerative Disease, New York Genome Center, New York, NY, USA; Department of Pathology and Cell Biology, Columbia University Medical Center, New York, NY, USA; Department of Neuroscience, Columbia University Medical Center, New York, NY, USA; The Carol and Gene Ludwig Center for Research on Neurodegeneration, Columbia University, New York, NY, USA

**Author notes:** <u>Corresponding author:</u> Marta Olah, PhD.

## Abstract

Development of therapeutic approaches that target specific microglia responses in amyotrophic lateral sclerosis (ALS) is crucial due to the involvement of microglia in ALS progression. Our study identifies the predominant microglia subset in human ALS primary motor cortex and spinal cord as an undifferentiated phenotype with dysregulated respiratory electron transport. Moreover, we find that the interferon response microglia subset is enriched in donors with aggressive disease progression, while a previously described potentially protective microglia phenotype is depleted in ALS. Additionally, we observe an enrichment of non-microglial immune cell, mainly NK/T cells, in ALS central nervous system, primarily in the spinal cord. These findings pave the way for the development of microglia subset-specific therapeutic interventions to slow or even stop ALS progression.

## Introduction

Amyotrophic lateral sclerosis (ALS) is a rapidly progressing neurodegenerative disease characterized by the gradual loss of upper and lower motor neurons, leading to a decline in motor function^1^. The median survival time from onset of symptoms is typically only 3-5 years, depending on specific subtype of the disease^2^. Currently, there are no disease-modifying treatments available for ALS, making it uniformly fatal.

One of the most prominent neuropathological features of ALS, in addition to motor neuron degeneration, is neuroinflammation. Human positron emission tomography (PET) imaging studies using TSPO radioligands have revealed that neuroinflammation can be detected early in the course of ALS and persists up until the later stage of the disease^3^. Neuroinflammation is associated with the activation of microglia, which are the resident innate immune cells of the central nervous system (CNS). Several histopathological studies using post-mortem brain and spinal cord specimens have demonstrated microglial proliferation and activation in ALS, with the severity of microglia-associated pathology closely correlating with disease progression^4,5^. While these studies have provided valuable insights, they mainly rely on morphological analysis or general markers for assessing microglial activation. As a result, our understanding of the complex molecular mechanisms underlying microglial involvement in ALS pathobiology remains limited. This knowledge gap is further emphasized by the failure of clinical trials that attempted to block general neuroinflammation in altering disease progression. Surprisingly, therapies targeting neuroinflammation that demonstrated significant benefits in pre-clinical rodent models of ALS have not translated into successful outcomes in human clinical trials. This disparity highlights the importance of studying the identity and role of microglia specifically in humans to gain a deeper understanding of ALS pathogenesis.

In our recent work, we made significant strides in this area by identifying nine human microglial subsets across different donor ages using single-cell RNA-sequencing^6^. Building upon this knowledge, we expanded our investigations to ALS donor samples using the same experimental and analytical pipeline. By comparing the results to our previous study, we successfully identified microglial and non-microglial immune cell subsets that are enriched in ALS. Furthermore, we explored the functional annotation and transcriptional regulatory networks of these ALS-associated microglia phenotypes, validating their relevance for ALS disease pathogenesis across independent datasets. Thus, our study provides a comprehensive account of the human microglial subsets associated with ALS, identified through unbiased transcriptomic approaches. Our findings will pave the way for developing therapeutic interventions that target specific microglia subsets and their unique pathways in the context of ALS.

## Results

### Establishing a microglia population structure in human ALS brain and spinal cord

To profile the microglial phenotypes present in the CNS of sporadic ALS in an unbiased fashion, we utilized our optimized pipeline for the cold, non-enzymatic isolation and single-cell RNA-sequencing (scRNA-seq) of live microglia and non-microglial immune cells (all CD45^+^) from autopsy brain and spinal cord tissue^7^. This approach has been shown to be superior to other single-cell and single-nucleus RNA-sequencing approaches in terms of capturing the full extent of microglia phenotypes without inducing a stress response signature^8^. We sampled multiple CNS regions from sporadic ALS donors of both sexes: prefrontal cortex (BA9), primary motor cortex (BA4), motor nucleus of the facial nerve, and lumbar spinal cord (**Figure 1A**). Three of donors carried a mutations in the SOD1 gene and one of the donors C9orf72 repeat expansion carrier (Table S1). After quality control (see Table S1), we obtained >51,000 single-cell transcriptomes of microglia and non-microglial immune cells. To facilitate comparison to the microglia phenotypes that we recently identified in aged and young donors^6^, we applied a label transfer to assign the ALS-associated immune cells to the clusters that we had defined previously (microglial clusters 1-9 and non-microglial clusters 10-14)(**Figure 1B** and **Figure S1A** and **S1B**). Each microglial cluster is marked by distinct genes that robustly distinguish them from one another (**Figure 1C**, **Table S2**). The identified microglial subsets are also characterized by non-overlapping functional annotations (**Figure 1D**) and reside along divergent state transition trajectories (**Figure 1D** and **1E**), highlighting their unique, non-overlapping identities.

**Figure 1.**
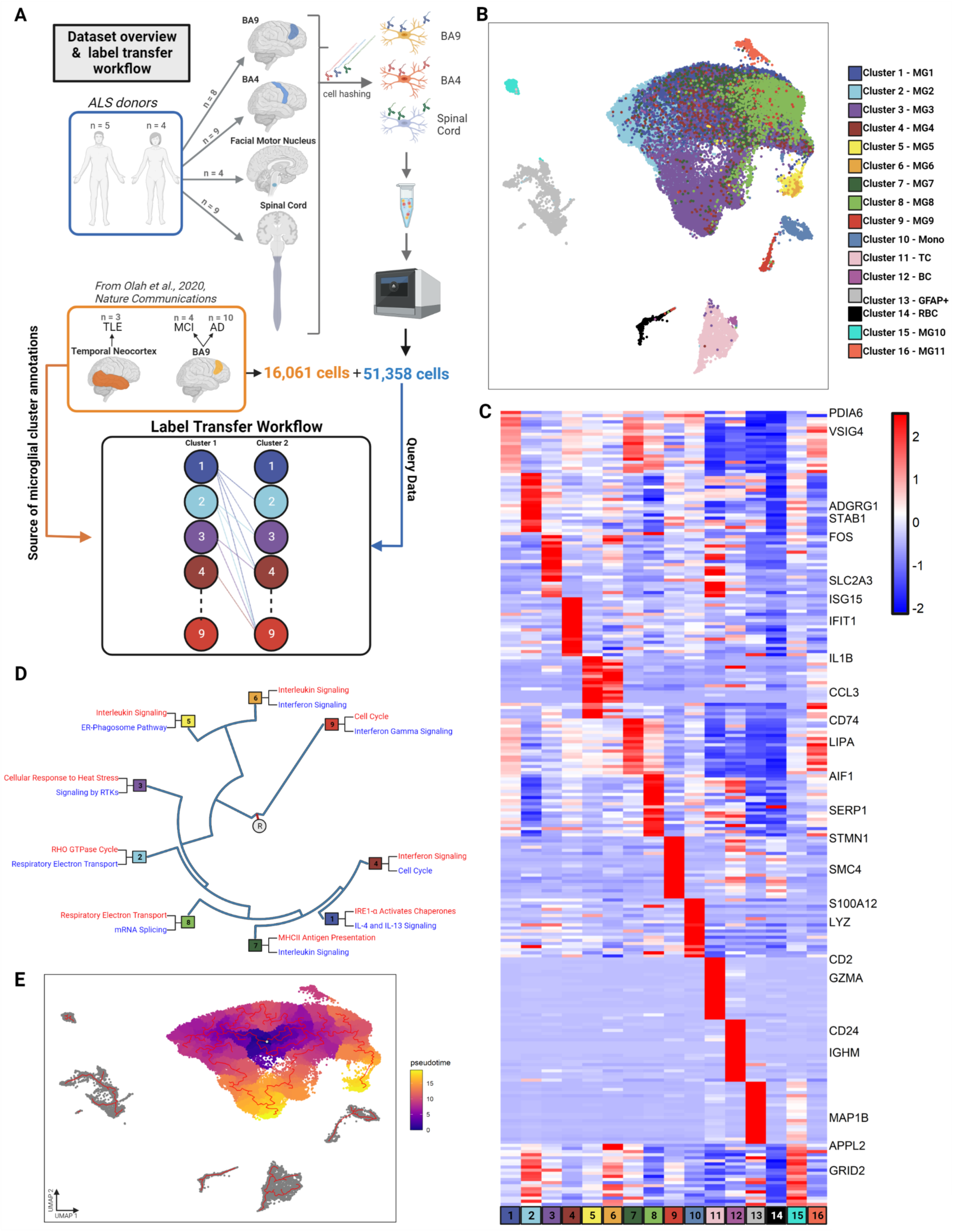
The population structure of microglia in human ALS brain and spinal cord. We used label transfer to explore the microglia (and other immune cell) phenotypes present in the central nervous system (CNS) of ALS donors. **(A) Sample collection and label transfer workflow.** Multiple CNS regions were sampled from 9 ALS donors with similar representation of both sexes. Single cell RNA sequencing data of live CD45+ immune cells was generated using the 10x Chromium platform. The ALS data was mapped onto our previously published microglia population structure by utilizing a pairwise machine learning approach with random forest classifiers and consensus voting to identify final labels. **(B) UMAP projection of the merged dataset.** The merged dataset is plotted on the first two UMAP components following Harmony batch correction. Each dot is a single cell. Microglia cluster 15 (MG10) had no unique gene set, while cluster 16 (MG11) was present only in one donor, accordingly they were not included in downstream analysis. **(C) ALS microglia subsets have unique marker gene sets.** Heatmap representing Z-scored expression data. Each column is a single cluster and each row is a single gene. **(D) Microglial subsets present in ALS have distinctive functional annotations.** Hierarchical dendogram demonstrating the functional landscape of microglia subsets. Similarity of subsets was calculated using Euclidean distance across the average expression profiles of each cell subtype. R denotes the root node. Top up– and down-regulated terms were selected from Reactome pathway annotation to highlight unique aspects of each microglial subsets. Terms in red are upregulated in a given cluster while terms in blue are downregulated. **(E) Microglia subsets in ALS reside along divergent state transition trajectories.** A pseudotime trajectory was built with monocle3, setting the root point in the middle of cluster 1. The trajectory is highlighted in red, showing the shifts seen across different aspects of the microglial cloud, including different pseudotime endpoints. Abbreviations: BA Brodmann area, ALS amyotrophic lateral sclerosis, UMAP uniform manifold expression and projection, R root node.

### ALS is characterized by robust, region-specific shifts in microglial subtype prevalence

We first aimed to identify the global changes in microglia subset abundance in ALS, using our published dataset of young and old non-ALS microglia^6^ as a reference. We observed robust, statistically significant shifts in the population structure of microglia in ALS (**Figure 2A**, **Table S3**). These changes are dominated by an inverse relationship between the two homeostatic microglia subsets MG1 and MG2. While MG1 is the most abundant microglial cluster in non-ALS samples, MG2 is the predominant microglia cluster in ALS. Furthermore, ALS samples also had higher representation of stress-related microglia cluster MG3, and lower proportions of the MHC-pathway enriched microglial cluster MG7. The higher prevalence of MG2 in ALS was prominent in all brain regions examined as well as in the spinal cord (**Figure 2B**, **Figure S2B**). In contrast, the enrichment of MG3 was more dramatic in ALS spinal cord samples and less so in ALS brain samples (**Figure 2B**, **Figure S2C**), while the microglia subset associated with respiratory electron transport (MG8) showed a trend to be enriched in ALS, especially in the subcortical samples. The non-microglial clusters 11 (**Figure 2B**), identified as T cells, were also found to be significantly enriched in ALS.

**Figure 2.**
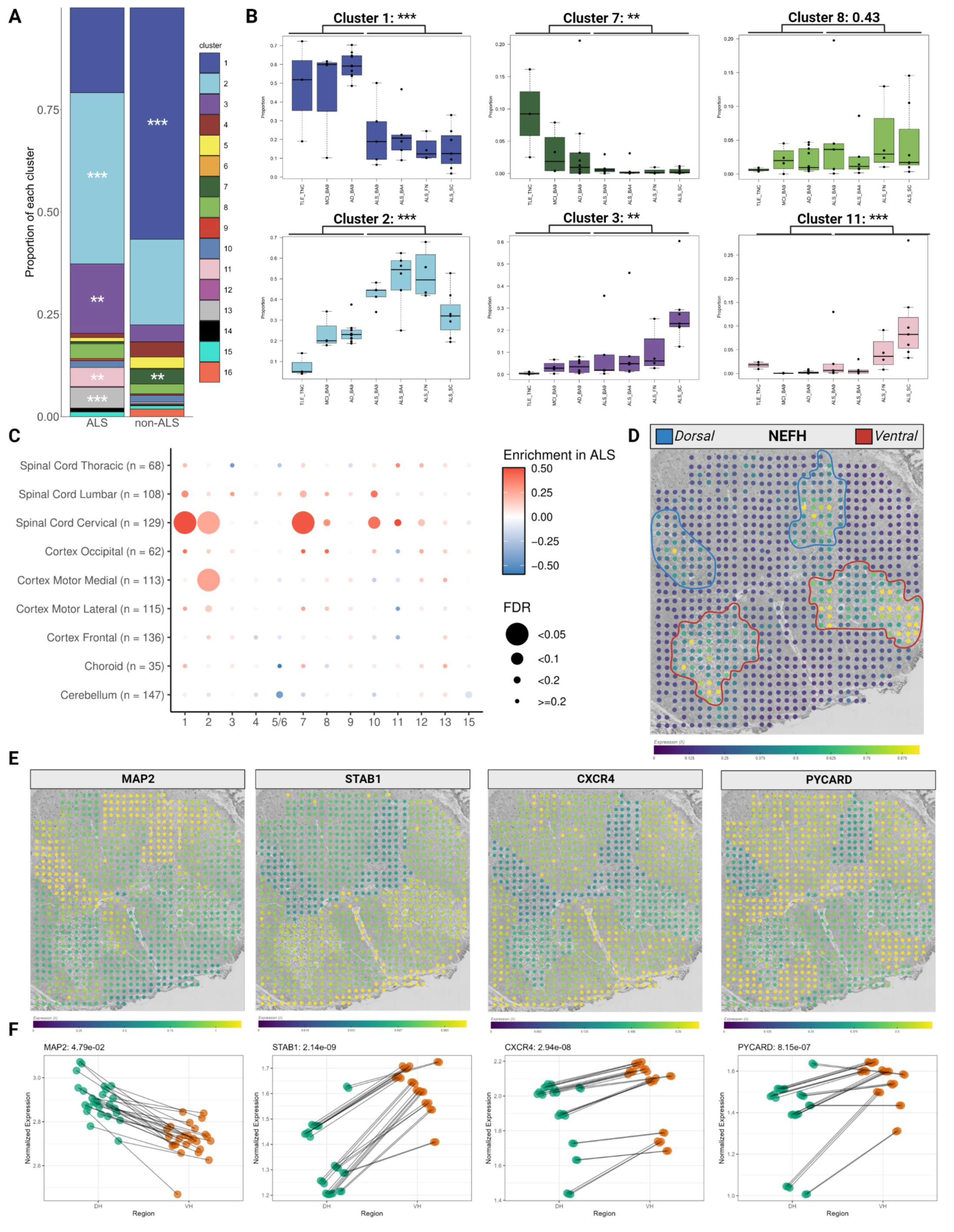
ALS induces robust, region specific shifts in microglial subtype prevalence. (A) Global changes in microglia subset abundance in ALS. Stacked bar chart showing the overall changes in immune cell abundance. Each bar shows the proportion of the different microglial and non-microglia immune cell subsets in each condition: non-ALS and ALS. Asterisks denote significant difference between conditions, using the Wilcoxon rank-sum test and BH correction to determine significance. *** <0.005, ** <0.01, * <0.05 **(B) Region specific changes in microglia cluster abundance in ALS.** Boxplots showing the distribution of individual cluster abundances across disease-region pairings. Representative clusters with significant, region specific changes are shown. Boxplots denote the 25^th^ percentile, median, and 75^th^ percentile, with whiskers capturing 1.5 IQR in both directions. **(C) Orthogonal validation in an independent bulk RNA-seq dataset confirms consistent association of microglia Cluster 2 signature with ALS.** Using a separate bulk RNA-seq cohort of 170 samples from ALS patients and non-ALS neurological disease patients or control patients, signatures of the top 20 genes per cluster were used to delineate the enrichment of different cluster signatures in ALS versus non-ALS samples. Notably, cluster 2 is the only cluster that shows significant enrichment in ALS in multiple regions, while cluster 1 and 7 enrichment likely capture the increase in overall microglia numbers in ALS. **(D) Orthogonal validation in an independent spatial transcriptomic dataset confirms the upregulation of ALS associated microglial subset marker genes at the anatomical sites of motor neuron death in ALS.** Spatial transcriptomic data was repurposed from Maniatas et al. A representative image from the slide viewer at https://als-st.nygenome.org/ is shown, displaying the neuronal gene NEFH, which is primarily found in the dorsal and ventral horns. Color bar is expression lambda calculated by the Splotch model. Dorsal and Ventral horns are demarcated in blue and red respectively. Representative images in following panels are from the same section. **(E)-(F) Marker genes of clusters 2 and 8 follow inverse patterns of upregulation in comparison to MAP2 in the dorsal and ventral horns of ALS patients.** Representative images for each gene as in (D) are shown in (E). In (F), dotplots compare the summed score of predicted counts for a given gene in all spots in the dorsal horn (green) to an identical score for that gene summed from all spots in the ventral horn (green) across a subset of donors with strong MAP2 detection. Testing for significance of differences between regions for each gene was conducted with Welch’s t-test and the Holm-Bonferroni correction. STAB1 is a defining gene for cluster 2, CXCR4 is a marker for cluster 3, and PYCARD is a defining gene for cluster 8. Abbreviations: BA Brodmann area, ALS amyotrophic lateral sclerosis, AD Alzheimer’s disease, MCI mild cognitive impairment, TLE temporal lobe epilepsy, ALS amyotrophic lateral sclerosis, BA Brodmann area, TNC temporal neocortex, SC spinal cord, SN substantia nigra, FN facial nucleus, IQR interquartile range, FDR false discover rate, DH dorsal horn, VH ventral horn.

We next performed orthogonal validation of our findings in an independent bulk RNA-seq dataset consisting of 170 samples from ALS patients, non-ALS neurological disease patients, and control donors^9^. Signatures of the top 20 genes per microglial cluster (**Table S4**) were used to delineate the enrichment of different cluster signatures in ALS versus non-ALS samples. This investigation confirmed that MG2-enriched genes had significantly higher expression in ALS motor cortex and cervical spinal cord when compared to controls (**Figure 2C**). Interestingly, MG1– and MG7-enriched genes also showed higher expression in the bulk RNA-sequencing dataset, but not in our scRNA-seq dataset. This may be because signature gene sets of MG1 and MG7 are particularly enriched in pan-microglial marker genes expressed by all microglia (**Figure S3**). Consequently, the enrichment of the MG1 and MG7 signatures in the bulk RNA-seq data likely reflects the expansion of the microglial population in the cervical spinal cord of ALS patients; a phenomenon that has been documented previously^10^. Interestingly, relative abundance of microglia (detected in the bulk RNA-seq dataset through the signatures of MG1 and MG7) positively correlated with family history of ALS as well as disease duration (**Figure S4B** and **S4D**), but not with sex, C9orf72 repeat expansion status, age at death, or age at onset (**Figure S4A**, **S4C**, **S4E** and **S4F**, respectively). MG2 was significantly positively associated with a positive family history of ALS, as were MG8 and MG9 and the monocytic cluster 10 (**Figure S4B**). Intriguingly, MG8 was negatively associated with disease duration along with MG4 (**Figure S4D**) and was significantly enriched in donors who had rapidly progressing disease (disease duration following diagnosis less then 2 years).

To further validate our scRNA-seq findings, we examined the spatial expression pattern of microglia subset-specific genes in an independent spatial transcriptomic dataset of ALS spinal cord^11^. Specifically, we investigated the differences in the expression levels of microglia cluster markers between the dorsal horn, which harbors intact neurons, and the ventral horn of the spinal cord, the site of motor neuron cell death. This dataset reliably showed reduced gene expression of *MAP2*, a neuronal marker, in the ventral horn of ALS spinal cord when compared to the dorsal horn (**Figure 2D, 2E** and **2F**, **Table S5**), consistent with extensive motor neuron demise in this area. Importantly, marker genes of MG2 (*STAB1*), MG3 (*CXCR4*) and MG8 (*PYCARD*) were inversely correlated with MAP2 expression in the dorsal and ventral horns of ALS spinal cord (**Figure 2E** and **2F**), suggesting their association with ongoing motor neuron death in the ventral horn of ALS spinal cord. Patterns of expression for other microglial subset-specific genes in the spatial transcriptomic dataset are shown in **Figure S4G**.

### ALS results in functional changes in microglia subsets

A central question in the analysis of microglial single-cell transcriptomics signatures is the degree to which the identified subsets are functionally distinct. We first established the non-overlapping functional identity of our microglial subsets using their unique signature gene sets and the REACTOME pathway analysis tool (**Figure 1D**, and **Figures S5A** and **S5B**, **Table S6**). We have previously reported that MG1 and MG2 are both a homeostatic/undifferentiated microglia subset^6^. In the combined dataset they maintained this annotation, with additional enrichment of ‘IRE1-a activated chaperons’ (MG1) and ‘RHO GTPase cycle’ (MG2). Similarly, the other microglia subsets also maintained their top functional annotation in the combined dataset: ‘Cellular response to heat stress’ for MG3, ‘Interferon Signaling’ for MG4, ‘Interleukin Signaling’ for MG5 & MG6, ‘MHCII Antigen Presentation’ for MG7, ‘Respiratory Electron Transport’ for MG8 and ‘Cell Cycle’ for MG9 (**Figure 1D** and **Figure S5A** and **S5B**).

To better understand microglial phenotypic changes in ALS, we used REACTOME for functional annotation^12^. First, we investigated the within-cluster transcriptional changes between ALS and non-ALS samples (**Figure 3A, Table S7**). The REACTOME terms that were upregulated within each cluster included pathways such as: ‘Chaperone mediated autophagy’ (upregulated in MG3 and MG9), ‘Role of phospholipids in phagocytosis’ (upregulated in MG1, MG4 and MG7), ‘Integration of energy metabolism’ (MG1, MG4, MG7 and MG8) and ‘G alpha (i) signaling events’ (upregulated in MG8) (**Figure 3A**). Most of these terms were upregulated in more than one microglia subset (e.g. ‘Interleukin-4 and Interleukin-13 signaling’ in MG1, MG3, MG4 and MG8), suggesting more global phenotypic changes in response to the ALS microenvironment. However, no responses were common to every cluster, suggesting that even in the face of a relatively similar microenvironment, the different microglial clusters may have different adaptive responses. Similarly, among the downregulated pathways, ‘Neutrophil degranulation’ was shared between 6 independent microglia clusters (MG1, MG2, MG3, MG5, MG6 and MG9), while others were shared between three microglia subsets (e.g. ‘Interferon gamma signaling’ in MG2, MG6 and MG9) or two subsets (e.g. ‘NGF-stimulated transcription’ in MG1 and MG5). On the other hand, downregulation of the ‘Respiratory electron transport chain’ pathway was only observed in MG2 while ‘Regulation of actin dynamics for phagocytic cup formation’ was specific to MG3 (**Figure 3B**). These results suggest microglia subset-specific metabolic and functional shifts that occur in the ALS microenvironment.

**Figure 3.**
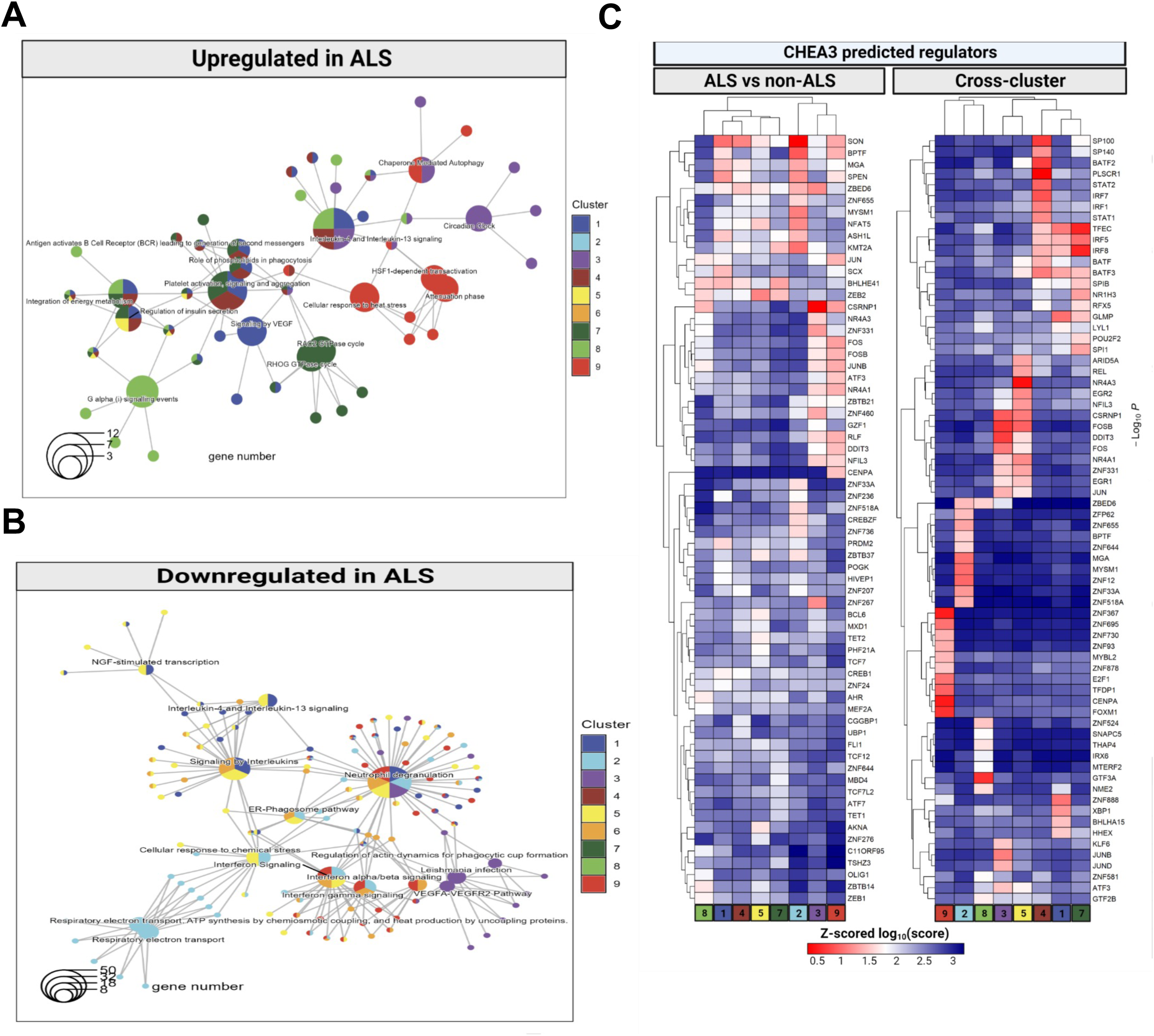
ALS results in functional changes in microglia subsets. Annotating between subset and within-subset shifts in ALS microglia at the level of transcriptome, proteome, and epigenome. **(A)-(B) Annotation of ALS associated transcriptional changes within each microglia cluster reveals functional shifts in ALS microglia.** REACTOME pathway analysis of differentially expressed genes within each microglia cluster between ALS and non-ALS. Results are displayed as a connectivity plot, where central nodes represent REACTOME pathways, while terminal nodes represent genes associated with those REACTOME terms. Central nodes are colored based on the enrichment of the term in a given cluster, while terminal nodes are colored based on the presence of the gene in the differentially expressed gene list for the ALS vs. non-ALS comparison for a cluster. Genes/terms upregulated in ALS are shown in (A) and genes/terms downregulated in ALS are shown in (B). Please note that many of the pathways are differentially expressed in more than one microglia cluster. **(C) Identification of differential and shared transcriptional regulators in cross-cluster and within-cluster cross-disease comparisons.** Both heatmaps show Z-scored log-normalized scores from CHEA3. On the left panel, genes used for regulator calculation were the top 50 genes derived from within-cluster across-disease differential expression for each cluster. On the right panel, genes used as input were selected top 20 marker genes per cluster. Rows and columns are clustered hierarchically by absolute linkage. Each row shows the scores for a single regulator across all microglia clusters (columns).

Next, we aimed to identify the regulators of gene expression governing microglial subset-specific and ALS-induced transcriptional programs. Using ChEA3^13^ we first predicted transcriptional regulators for the individual microglia subsets (right panel of **Figure 3C**, **Table S8**). As proof-of-concept, MG4, which showed strong enrichment in the ‘Interferon Signaling’ pathway, was predicted to be regulated by the transcription factors STAT1, STAT2, IRF1 and IRF7, while the stress-associated MG3 showed regulation by JUNB and JUND. ZNF888 seems to be a specific regulator of the MG1 transcriptional identity, BPTF, MYSM1 and MGA were specific to MG2, NR4A3 to MG5, GTF3A appeared to be specific to MG8, while E2F1 and CENPA target genes were enriched in MG9. Many of the transcriptional regulators of MG7 are shared with other clusters, especially with clusters MG1 and MG4, suggesting shared ontology for these microglia subsets (**Figure 1D**). In contrast, most of the ALS-associated gene expression changes within clusters were regulated by SON, a member of the nuclear speckle pre-mRNA processing machinery^14^ (left panel of **Figure 3C**, **Table S9**). One exception was MG8, which had very few transcriptional regulators that were shared with other clusters in this comparison (**Figure 3C**, left panel).

### Independent clustering of non-microglial immune cells identifies shifts in myeloid and adaptive immune cell populations in ALS

Since our dataset included other non-microglial immune cells (all CD45^+^ cells) isolated from CNS tissue, we performed additional de novo clustering exclusively on these cells to better understand their identity and relevance in ALS. We identified 19 non-microglial immune cell clusters that could be broadly classified as T cells, NK cells, macrophages, monocytes, B cells, and dendritic cells (**Figure 4A**, **Table S10**). These peripheral immune cells profiled from the brain parenchyma had non-overlapping signature gene sets (**Figure 4B**, **Table S11**) and showed divergent enrichment in ALS (**Figure 4C-H**, **Table S12**). Interestingly, the macrophage cluster C2 was depleted from most ALS CNS regions, except for BA4, while the stressed T cells (C3), T cells (C4), maturing NK cells (C5), NK cells (C8) and dendritic cells clusters (C10) were robustly enriched in ALS samples (**Figure 4C-H**). CD8+ (C7) and CD4+ T cells (C9) were also enriched in ALS when compared to Alzheimer’s’ disease (AD) and mild cognitive impairment (MCI) samples, but not when compared to samples from young donors (**Figure S6**). These findings were further corroborated by mapping the non-microglia immune cells to the Azimuth reference atlas^17^ and examining the relative abundance of the different non-microglial immune cell subsets in our dataset (**Figure S7A-7H** and **Figure S8**, **Table S13**). Similar to the de novo clustering, CD4 positive central memory T cells (T_CM_s) and CD8 T_CM_s identified through this reference mapping approach were also relatively enriched in ALS samples, although a substantial number of CD4 T_CM_s were observed in TLE samples as well (**Figure S7C** and **S7E**). The reference mapping approach also revealed the enrichment of _γδ_T cells in ALS samples as well as the depletion of plasmablasts in ALS (**Figure S7D** and **S7G**, respectively).

**Figure 4.**
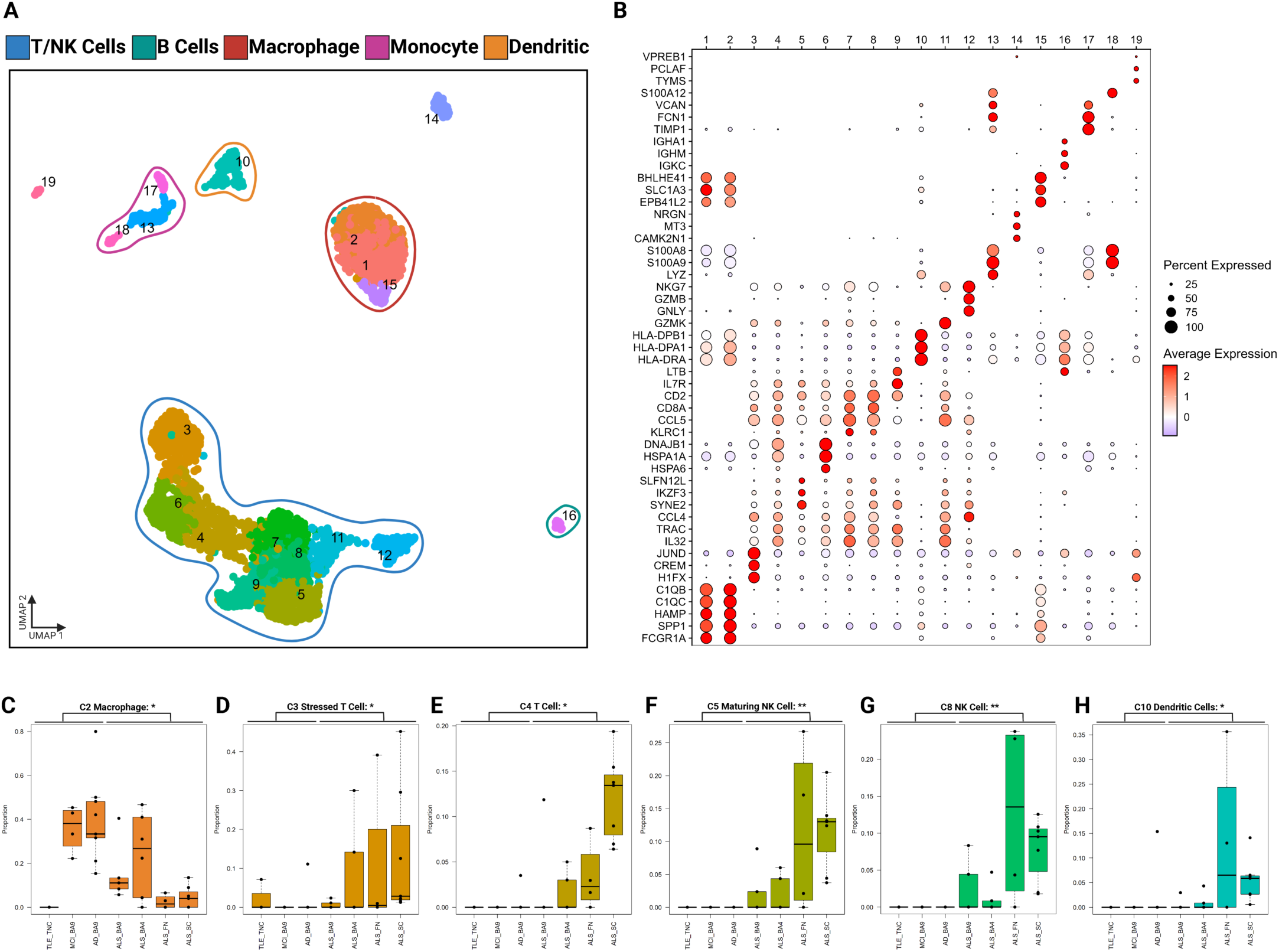
Independent clustering of non-microglial immune cells identifies shifts in myeloid and adaptive immune cell populations in ALS. (A) Independent clustering of non-microglial immune cells. Non-microglial immune cells were isolated in silico and separately processed. Optimized clustering resolution was chosen with ChooseR. Clusters with greater than 10 cells are plotted on the UMAP. **(B) Selected marker genes identify diverse immune populations.** Each column represents one of the clusters, and each row represents the z-scored expression of a given gene. The plot is colored according to level of expression and the size of each circle represents the percentage of cells in each cluster that express the gene. **(C)-(H) Annotation identifies enrichment of T and NK cells in ALS, as well as enrichment of dendritic cells and depletion of macrophages.** Boxplots denote the 25^th^ percentile, median, and 75^th^ percentile, with whiskers capturing 1.5 IQR in both directions. Asterisks denote significance of difference between conditions, using the Wilcoxon rank-sum test and BH correction to determine significance. * < 0.05. Abbreviation: ALS amyotrophic lateral sclerosis, NK natural killer.

## Discussion

In this study, we present a comprehensive assessment of human microglia phenotypes in ALS, using scRNA-seq of live microglia. Importantly, we document significant deviation from the observed changes of the microglia population structure in mouse models of this disease. To our surprise, MG7, the human microglia cluster that most closely resembled the murine Disease Associated Microglia (DAM) phenotype^6^ that was found to be enriched at every symptomatic stage in mouse SOD1 model^20,21^, was significantly depleted in human ALS, similarly to what we observed in AD^6^. Our findings suggest that the depletion of the putatively protective MG7 microglia phenotype might be a common feature of late-stage neurological diseases in humans and highlights the limitations of mouse models to study microglia phenotypes associated with age-related neurodegeneration due to species differences in microglia aging ^22,23,24^. Additionally, here we document a robust, region-specific reorganization of microglia population structure in this disease. In contrast to AD, MCI, and temporal lobe epilepsy (TLE), the dominant microglia phenotype in ALS brain and spinal cord was MG2, a homeostatic/undifferentiated microglia subset characterized by a profound restructuring of the oxidative phosphorylation machinery, through the downregulation of several electron transport chain subunits. Using an independent bulk tissue transcriptomic dataset^9,10^ we confirmed the significant enrichment of the MG2 expression signature in ALS motor cortex and spinal cord, when compared to healthy controls. The other prominent feature of ALS spinal cord was the strong enrichment of the stress– and early-response associated microglia cluster, MG3, in this region. In an independent spatial transcriptomic dataset^11^ gene signatures of both MG2 and MG3 showed an inverse relationship with MAP2, a neuronal marker depleted from ALS spinal cord ventral horn, suggesting that these subsets may localize specifically to areas of motor neuron demise. While MG2 and MG3 signatures did not show an association with disease duration, two smaller microglia subsets, MG4 and MG8, were found to be significantly enriched in the spinal cord of donors with aggressive disease progression. Interferon response, the functional annotation of the MG4 microglia cluster, has been previously shown to be the dominant pathogenic feature of C9orf72-/-myeloid cells and microglia^25,26^. We also found MG8 to be enriched in ALS brain and spinal cord in our dataset. This small microglia cluster was a mirror image of MG2 in terms of functional annotation (upregulated genes in MG2 were downregulated in MG8, and vice versa). Intriguingly, signature markers of the ALS enriched protein co-expression module^27^ were expressed at the highest level in MG8 in our dataset, as were many of the known interacting partners of TDP-43^28^, including SOD1 and CHCHD10; both implicated in familial forms of ALS^29,30^ (**Figure S8**). Furthermore, the novel ALS cerebrospinal fluid biomarker candidates CAPG^31^ was most prominently expressed in MG8 among the different microglia clusters (**Figure S8**). Importantly, none of these transcriptomic microglia phenotypes could be deduced from bulk tissue RNA sequencing data^9,10,32^, nor could they be identified in a single-nucleus RNA sequencing dataset^33^, confirming the importance of ex-vivo (whole cell) microglia profiling for the study of transcriptional heterogeneity in this cell type^34^.

Additionally, an examination of non-microglial immune cells in ALS revealed a preponderance of T cells, as well as relative enrichment of dendritic cells and NK cells in the ALS spinal cord. Interestingly, annotation of this subset of our dataset with the Azimuth PBMC reference^17^ highlights enrichment of γδT cells in ALS, which have been reported to be increased in the peripheral blood of ALS patients, but have not been explored yet in the CNS^35^. Our findings open novel avenues for further investigations into the role of the different infiltrating peripheral immune cells in the ALS spinal to understand whether they are contributing to the pathobiology directly or through their interaction with local microglia.

Despite the challenges associated with ex-vivo studies of human microglia in ALS, we believe that the approach described here is crucial to understanding of their role in ALS disease pathobiology. The ALS-enriched human microglia phenotypes presented here have not been recapitulated in murine or in vitro model systems yet. Accordingly, mechanistic studies of their role in ALS models is challenging at this moment. Nonetheless, our data set will enable the establishment of translational studies that aim to generate novel in vitro and preclinical model systems in which drug screens targeting specific microglia subsets can be performed to halt or slow down ALS disease progression.

## Acknowledgements

We are grateful to the donors and their families who made this study possible with the donation of brain and spinal cord tissue at autopsy. Some of the experiments (cell sorting) reported in this publication were performed in the CCTI Flow Cytometry Core, supported in part by the Office of the Director, National Institutes of Health under awards S10OD020056. Single-cell RNA sequencing was performed at the Columbia University Medical Center Genome Center Single Cell Analysis Core, which is funded in part through the NIH/NCI Cancer Center Support Grant P30CA013696 as well as by the National Center for Advancing Translational Sciences/NIH through grant number UL1TR001873. SP is supported by NS117583 and NS107442. Data acquisition and analysis were also supported by a Chan-Zuckerberg Neurodegeneration Challenge Network grant (CS-02018-191971) and NIH RF1 AG057473, P30 AG066462, and U01 AG061356.

## Author contributions

Conceptualization: J.F.T., M.O.; data curation: J.F.T., M.F., S.M., A.D., V.M., M.O.; formal analysis: J.F.T., M.F., V.M., M.O.; funding acquisition: N.S., H.P., P.L.D., S.P., M.O.; investigation: J.F.T., A.K., C.H., M.O.; methodology: J.F.T., A.K., C.H., M.O.; project administration: M.O.; resources: N.S., A.F.T., H.P., P.S., J.A.S., D.A.B., P.L.D.; software: J.F.T., V.M.; supervision: V.M., O.M.; validation: J.F.T., V.M., M.O.; visualization: J.F.T., M.F.; writing – original draft: J.F.T. & M.O.; writing – review & editing: all authors.

## Materials and methods

### Table of key resources

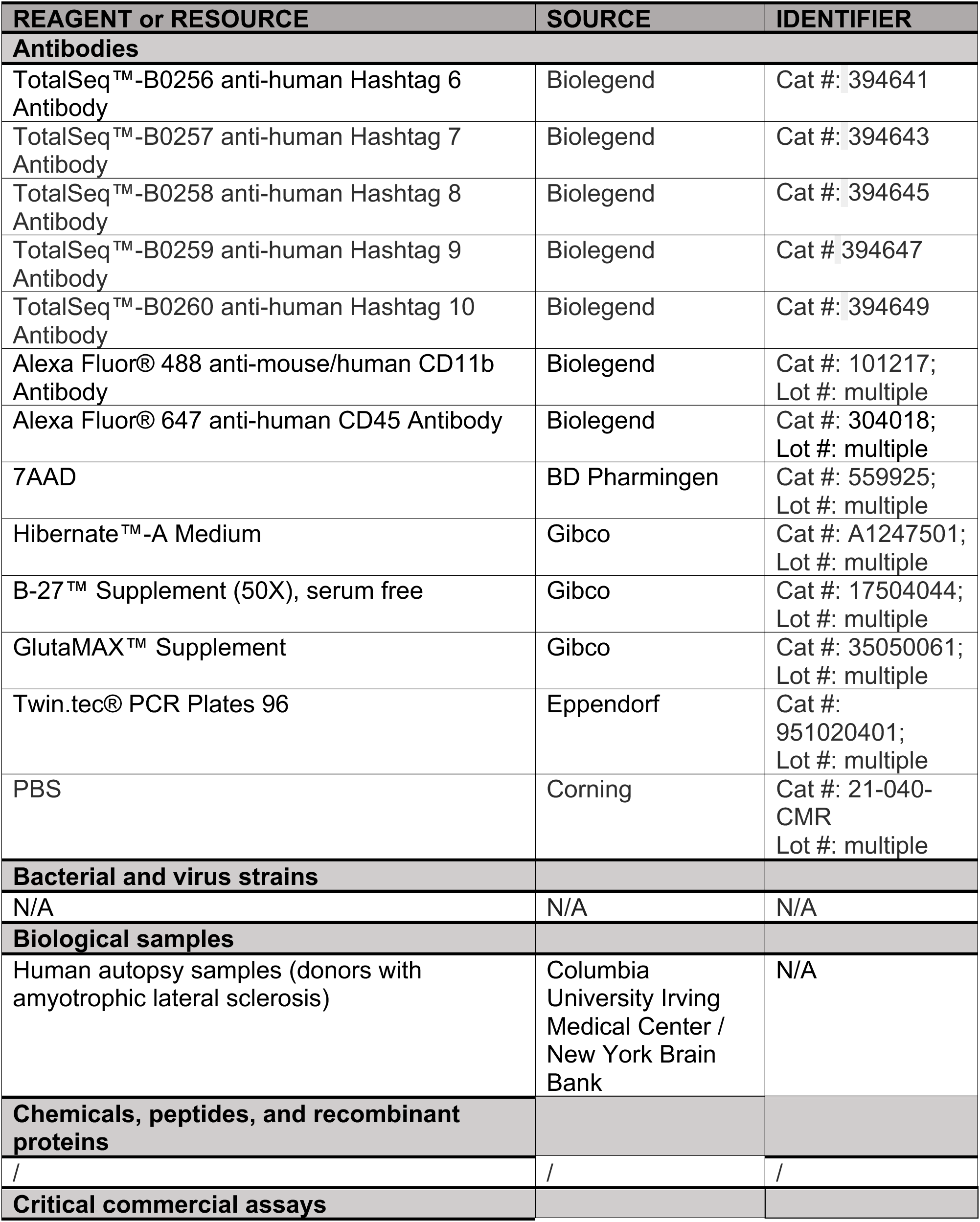
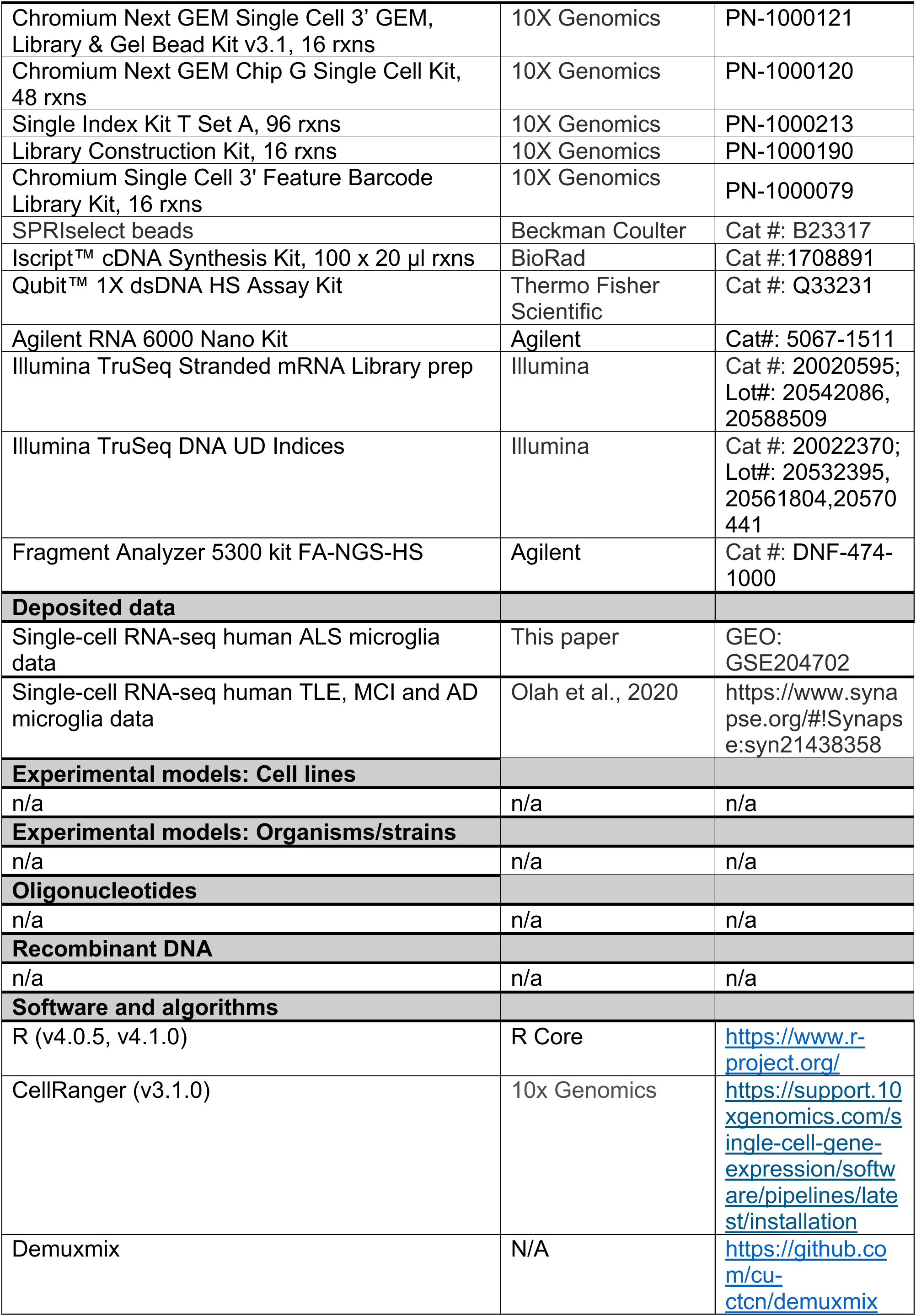
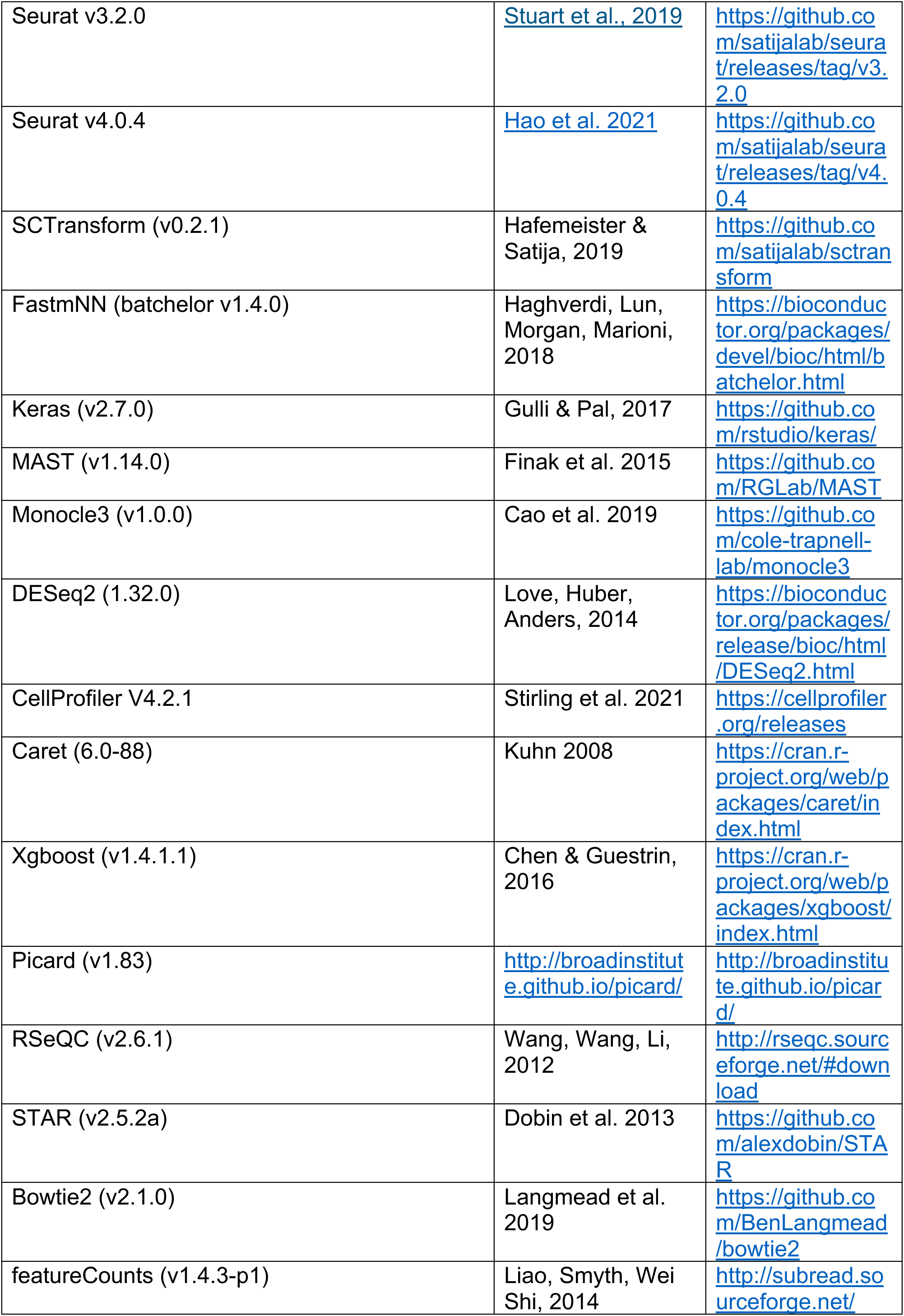
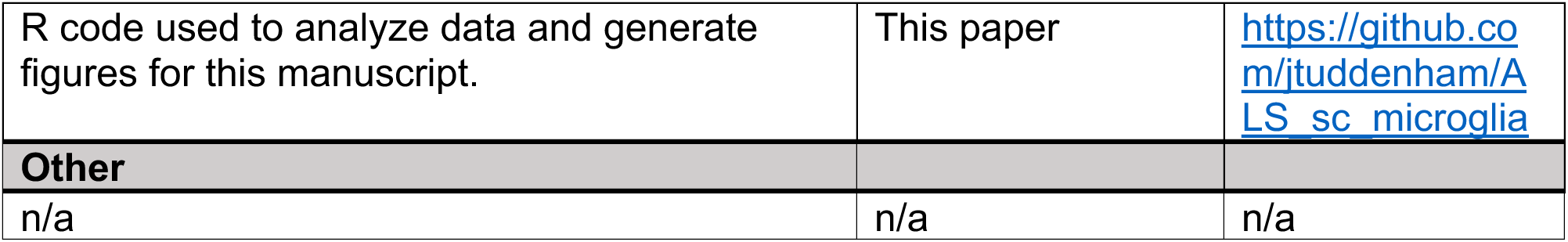

#### Source of central nervous system specimens

All brain specimens were obtained through informed consent and/or brain donation program at the Columbia University Medical Center (CUMC) / New York Brain Bank (NYBB). All procedures and research protocols were approved by the Institutional Review Board (IRB) of Columbia University Medical Center (protocol AAAR4962). For a detailed description of the brain regions sampled, clinical diagnosis, age and sex of the donors see **Table S1**. After weighing, the resected brain and spinal cord tissue was placed in ice-cold transportation medium (Hibernate-A medium (Gibco, A1247501) containing 1% B27 serum-free supplement (Gibco, 17504044) and 1% GlutaMax (Gibco, 35050061)) and transported from the autopsy suite at CUMC/NYBB to the wet lab at CUMC on ice (4°C) for immediate processing and live microglia isolation.

#### Microglia isolation, cell hashing and sorting

The isolation of microglia was performed according to our published protocol^4^, with minor modifications. In case of the cortical autopsy samples (BA9/46, BA4), the cortex (grey matter and the underlying white matter (subcortical white matter)) were dissected under a stereomicroscope. The subcortical white matter samples were not used in this study. The spinal cord sample (SC) was sampled at the level of lumbar section and included both white and grey matter. All steps of the protocol were performed on ice. The dissected tissue was placed in HBSS (Lonza, 10-508F) and weighed. Subsequently the tissue was homogenized in a 15 ml glass tissue grinder, 0.5 g at a time. The resulting homogenate was filtered through a 70 um filter and spun down at 300rcf for 10 minutes. The pellet was resuspended in 2 ml staining buffer (RPMI (Fisher, 72400120) containing 1% B27) per 0.5 g of initial tissue and incubated with anti-myelin magnetic beads (Miltenyi, 130-096-733) for 15 minutes according to the manufacturer’s specification. The homogenate was than washed once with staining buffer and the myelin was depleted using Miltenyi large separation columns (Miltenyi, 130-042-202). The cell suspension was spun down and was then incubated with anti-CD11b AlexaFluor488 (BioLegend, 301318) and anti-CD45 AlexaFluor647 (BioLegend, 304018) antibodies as well as 7AAD (BD Pharmingen, 559925) for 20 minutes on ice. Samples ALS1-ALS5 (**see Table S1**) were received and processed before the commercial availability of cell hashing antibodies. For samples ALS6-ALS9 the individual central nervous system regions were hashed using cell hashing antibodies along with anti-CD11b, anti-CD45 and 7AAD for 20 minutes on ice (for catalogue numbers of cell hashing antibodies see **Table S1**). Subsequently the cell suspension was washed twice with staining buffer, filtered through a 70 µm filter and the CD45+/7AAD-cells were sorted on a BD Influx cell sorter. Cells from each brain region were sorted in a separate A1 well of a 96 well PCR plate (Eppendorf, 951020401) containing 100 µl of PBS buffer with 0.3% BSA. For samples ALS1-ALS5 each sample/region was loaded independently on a 10x Chromium channel. For samples ALS6-ALS9 following sorting cells from different brain regions were combined and immediately submitted to single cell capture) 10× Chromium channel), reverse transcription and library construction on the 10x Chromium platform. All sorting was performed using a 100 µm nozzle. The sorting times varied according to the quality of the sample but was on average between 10 and 20 minutes per sample. The sorting speed was kept between 8000 – 10,000 events per second.

#### 10x Genomics Chromium single cell 3’ library construction

Cell capture, amplification and library construction on the 10x Genomics Chromium platform was performed according to the manufacturer’s publicly available protocol. Briefly, viability was assessed by trypan blue exclusion assay, and cell density was adjusted to 175 cells per μl. 7,000 cells were then loaded onto a single channel of a 10x Chromium chip for each sample. The 10x Genomics Chromium technology enables 3’ digital gene expression profiling of thousands of cells from a single sample by separately indexing each cell’s transcriptome. First, thousands of cells are partitioned into nanoliter-scale Gel Bead-In-EMulsions (GEMs). Within one GEM all generated cDNA share a common 10x barcode. Libraries were generated and sequenced from the cDNA, and the 10x barcodes were used to associate individual reads back to the individual partitions. To achieve single cell resolution, the cells were delivered at a limiting dilution. Upon dissolution of the Single Cell 3’ Gel Bead in a GEM, primers containing (i) an Illumina R1 sequence (read 1 sequencing primer), (ii) a 16 nucleotide 10x Barcode, (iii) a 10 nucleotide Unique Molecular Identifier (UMI), and (iv) a poly-dT primer sequence were released and mixed with cell lysate and Master Mix. Incubation of the GEMs then produced barcoded, full-length cDNA from poly-adenylated mRNA. After incubation, the GEMs were broken and the pooled fractions were recovered. Full-length, barcoded cDNA was then amplified by PCR to generate sufficient mass for library construction. Enzymatic fragmentation and size selection were used to optimize the cDNA amplicon size prior to library construction. R1 (read 1 primer sequence) were added to the molecules during GEM incubation. P5, P7, a sample index, and R2 (read 2 primer sequence) were added during library construction via end repair, A-tailing, adaptor ligation, and PCR. The final libraries contained the P5 and P7 primers used in Illumina bridge amplification. The described protocol produced Illumina-ready sequencing libraries. A Single Cell 3’ Library comprises standard Illumina paired-end constructs which begin and end with P5 and P7. The Single Cell 3’ 16 bp 10x Barcode and 10 bp UMI are encoded in Read 1, while Read 2 is used to sequence the cDNA fragment. Sample index sequences were incorporated as the i7 index read. Read 1 and Read 2 are standard Illumina sequencing primer sites used in paired-end sequencing. Sequencing the library produced a standard Illumina BCL data output folder. The BCL data includes the paired-end Read 1 (containing the 16 bp 10x Barcode and 10 bp UMI) and Read 2 and the sample index in the i7 index read.

#### Batch structure and sequencing

Tissue specimens were processed upon receipt. The different central nervous system regions from the same donor were processed parallel. Samples from donors ALS1-ALS5 were loaded 1 region per 10x Chromium channel. The different central nervous system region samples from donors ALS6-ALS9 were hashed, combined and loaded 1 donor per 10x Chromium channel. Since each central nervous system region was processed from any given donor was processed parallel on the same day, each sample constitutes one batch for microglia isolation, cell capture and library construction. All sequencing was performed on an Illumina NovaSeq6000 machine. For specifics on sequencing and QC metrics regarding the generated reads see **Table S1.**

#### Single-cell RNA-seq data processing, alignment, and hashtag deconvolution

The majority of our downstream analysis was conducted using the R programming language (v4.0.5 for harmonization and clustering, v4.1.0 for annotation and downstream visualization)^2^ and the RStudio^3^ integrated development environment. CellRanger V3.1.0 with default parameters was used to demultiplex and align our barcoded reads with the Ensembl transcriptome annotation (downloaded March 2019, GRCh38.91). A recent report^4^ suggested that filtering cells with greater than 10% mitochondrial reads is the preferred baseline for human tissue, and that for brain tissue a higher threshold may even be optimal. Thus, a mitochondrial percentage that was the higher of either 10% of reads or the 2 absolute deviations above the median for mitochondrial reads within the sample was chosen as a threshold. Cells below this threshold with between 500 and 10,000 UMIs were retained for downstream analysis. All ribosomal genes, mitochondrial genes, and pseudogenes were removed, as they interfered with the downstream differential gene expression. For samples where we used cell hashing to combine regions or subjects in a single sequencing run, droplets were demultiplexed using the following workflow. For each HTO, a mixture model with two components was fitted to the HTO counts using an EM algorithm. The component with the smaller mean (negative component) represents droplets that were not tagged with the HTO, whereas the component with the larger mean (positive components) represents droplets that were tagged. We then assign each droplet to either the negative or positive component based on its posterior probability. Droplets that were assigned to the negative component for all HTOs as well as multiplets were discarded. Singlets with high uncertainty, i.e. without confident assignment to either the negative or positive component, were discarded as well, leaving only high certainty singlets for downstream analysis. The method is implemented in the R package demuxmix which available on github: https://github.com/cu-ctcn/demuxmix. Some of our hashtag data had lower overall counts, and thus, the demuxmix model was unable to effectively segregate distributions for some hashtags in several samples. These samples were identified as having high percentages of negative/uncertain cells with demuxmix. In these cases, to try and recover cells for further analysis, the problematic hashtags were reclassified using one of two different algorithms, a demixing algorithm developed for MULTI-seq^5^ or HTOdemux from Seurat v3.2.0^6^. Hashtag classifications were merged, and doublet/negative/uncertain cell removal proceeded as before.

#### Label transfer to annotate ALS single-cell RNA-sequencing data

Cluster labels were assigned to the cells obtained from ALS donors, as follows:

##### Assigning non-microglial identities

1) Expression data for all cells from all donors were first normalized using the SCTransform algorithm in Seurat, then integrated using the Harmony R package with 30 PCs and theta=5 and each donor as a batch, and subsequently clustered using 15 Harmony dimensions using the standard Louvain algorithm in the Seurat package (version 4) with resolution = 0.5.
2) T-cells, B-cells, monocytes, red blood cells, and GFAP+ clusters were identified using a combination of known gene markers, as well as composition of clusters 10-14 in the cells from non-ALS donors, as described in Olah et al. (2020)
3) In this process, we also identified two additional clusters that were assigned cluster labels 15 and 16, since they did not correspond to any of the 14 clusters previously identified in Olah et al. (2020).
4) The remaining cells, which were putatively assigned microglial identity, were then assigned identities using a bootstrapped random forest approach, as described below.

##### Assigning microglial cluster identities

5) A training set of 200 cells randomly selected from each of the microglial clusters in Olah et al. (2020) was constructed. For clusters with fewer than 200 cells, sampling was done with replacement.
6) For each cluster (1-9), differential genes distinguishing that cluster from all other cells were identified using a Mann-Whitney test on the SCTransformed cells from the training set, with nominal p-value<0.01
7) All 9 gene sets from step 6 above were then combined (removing duplicates) to form the master gene set for random forest training.
8) A random forest classifier was then constructed on the training set using the combined gene set in step 7, and subsequently run on all of the ALS donor cells, none of which were included in the training set.
9) Steps 5-8 were run 20 times, with a different random seed each time.
10) Cells from ALS cells were then assigned to the most commonly classified cluster identity (plurality voting) over the 20 runs.

After assignment of both the non-microglial and microglial cells, the entire data set was visualized using the same Harmony-based integration described in step 1 above, followed by standard UMAP implementation in Seurat (v4) with default parameters and 15 Harmony dimensions.

The final cluster labels thus included the 9 microglial and 5 non-microglial clusters from Olah et al. (2020), which now included cells both from that original data set as well as the new cells from ALS donors, and 2 additional clusters that did not emerge in the clustering from our original data set.

#### 10x chemistry correction

Striking differences were observed in the distributions of UMI counts between 10X v2 and v3 chemistry. As this was driving differential clustering, count matrices from v3 samples were downsampled by 50% using the DropletUtils^7^ package in R to achieve comparable UMI distributions across the two technologies. This was done prior to downstream visualization of gene expression and computation of differentially expressed genes between ALS and non-ALS microglia.

#### Identification of cluster-defining gene sets

To identify cluster-defining gene sets, the *FindMarkers* function in Seurat was used to implement a pairwise testing approach. We prioritized differentially expressed genes that could best delineate a given cluster from each other cluster in our dataset. To do so, MAST^8^ was applied to normalized count data from the “RNA” assay of the Seurat object to find differentially expressed genes between every combination of pairs of clusters. Within each cluster, all the differentially expressed genes that were identified with this approach were filtered to only include those that were only found to be differentially expressed in one direction (either up or down). Any genes that were found to be upregulated in comparison to some clusters but downregulated in comparisons to other clusters or vice versa were removed from our downstream analysis. Furthermore, to ensure that the specific cluster-defining genes were prioritized, upregulated genes were ranked by the number of comparisons in which they were upregulated, and only those upregulated in 3 or more comparisons were used for downstream analyses. An identical process was applied for downregulated genes. Full marker gene lists are reported in **Table S2**.

#### Identification of differentially expressed genes between ALS and non-ALS microglia

To identify cluster-defining gene sets, each cluster was separated into ALS and non-ALS microglial cells, then the *FindMarkers* function in Seurat was used, leveraging MAST to identify differentially expressed genes between the ALS and non-ALS conditions. The downsampled (see **10x chemistry correction**) normalized count data was used for this purpose to mitigate technical bias.

#### Constructing phylogenetic tree of microglial subtypes

To construct a phylogenetic tree evaluating the similarity of different microglial subtypes, the *BuildClusterTree* function in Seurat was used to build a phylogenetic tree on a distance matrix of “average” cells for each of the identity classes computed from the first 30 principal components. Results were visualized with the ggtree^9^ package, as in **Figure 1D**.

#### Functional annotation of microglial clusters

To perform functional annotation of microglial clusters, the top 50 up-or down-regulated genes in the differentially expressed gene lists for each cluster were taken for analysis. Annotation of these gene lists was performed with Reactome^10^ pathway analysis using clusterProfiler^11^. For all functional analysis, the Benjamini-Hochberg correction^12^ was used to correct p-values for multiple testing. Corrected p-values below a threshold of 0.05 were chosen as significant for Reactome results. This type of annotation was also performed on the ALS vs. non-ALS differentially expressed gene lists for each microglial cluster, as well as in the non-microglial immune cell analysis.

#### Monocle3 pseudotime analysis

As an orthogonal method of evaluating the continuity of different microglial states in our cluster structure, the monocle3 algorithm was used to build a pseudotime trajectory across our dataset as shown in **Figure 1E**. Using the Seurat interface to monocle3 found in SeuratWrappers, the Seurat object was converted into a monocle data object, and a pseudotime trajectory was derived using the *learn_graph* function, retaining the final cell identity assignments from our original label transfer and clustering pipeline (see **Label transfer to annotate ALS single-cell RNA-sequencing data**).To establish an originating point, the pseudotime root was placed on the border of clusters 1 and 2, as these subtypes were described in our prior paper^13^ as being the two most prevalent microglial subtypes. Interestingly, this state was best captured by choosing cells with maximal *AVP* expression, a marker of hematopoietic stem cells^14^ that is frequently used to mark the root cells in hematopoietic pseudotime tracing.

#### Identifying differentially represented subsets between ALS and non-ALS samples

To compare the abundance of different subsets between ALS and non-ALS samples, proportions of different cell types or subtypes were aggregated at the level of donor-region pairings. For example, ALS donor 1 would be represented by ALS1-Spinal Cord, ALS1-BA9, and ALS1-BA4. From here, each donor-region is treated as one sample for the purposes of comparing proportions of specific subtypes between ALS and non-ALS samples. A Wilcoxon rank-sum test was used to compute the significance of differences in proportion of each subset between ALS and non-ALS regions, and BH correction was used to correct p-values for multiple testing.

#### Identifying differentially represented subsets between regions in ALS samples

Similar to above, samples were aggregated at the level of donor-regions, and examined for differences between different groups. In this case, the generalization of the Wilcoxon, the Kruskal-Wallis test, was used to assess the significance of differences in proportion between different regions in ALS, and the BH correction was used to account for multiple testing.

#### Annotating single-cell microglial signatures in bulk RNA sequencing data from ALS donors

Bulk RNA-seq BAM files of the Target ALS Foundation were downloaded from the New York Genome Center in September 2020. These were 1,063 BAM files of spinal cords and brains from 208 unique donors. BAM files were reverted to FASTQ files using the SamToFastq function of Picard tools (v2.17.4). RNA-seq reads were mapped onto the reference human genome GRCh38 using the STAR aligner (v2.5.3a) with 2-pass mapping mode. Gene expression levels were quantified using RSEM (v1.2.31) with the Ensembl human gene model (release 91). Following samples were excluded at the quality control stage: one of samples sequenced twice, samples that have >3% ribosomal RNA, >40% duplication rate, <4 RNA integrity number (RIN), and a sample whose library preparation method (manual vs. automated) was missing from metadata. Donors were retained if they were diagnosed as ALS or non-neurological control. Tissues that had less than 10 samples were also excluded. After these filtering steps, 913 samples from 170 unique donors were included for subsequent analysis. The voom function of the R package limma was used to compute log_2_ counts-per-million-mapped-reads (CPM) adjusted for RIN and library preparation methods. For each gene, log_2_CPM expression levels were further normalized into z-scores. Microglial signature scores were computed by averaging z-scores of log_2_CPM over member genes.

#### Computing pseudobulk PCA on single-cell microglial data

After downsampling to mitigate technical bias, gene expression was aggregated at the donor-region level (i.e. summing all counts for cells derived from a specific donor-region combination). Principal components analysis was computed on this pseudo-bulk expression data, and the first two components were plotted.

#### Analyzing ALS spinal cord spatial transcriptomic data

Initial analysis of the spatial transcriptomic data from Maniatis et al. 2019^15^ showed infrequent expression of microglial genes and cluster markers due to the sparsity of the data. As such, the posterior lambda values that result from the use of the Splotch model were obtained from the authors of the original paper and used for further analysis. These posterior values were converted to predicted counts with the formula:

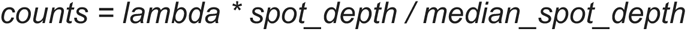

Where *median_spot_depth* was set to 1203. Samples with good quality *MAP2* signal were manually selected from the slide viewer at https://als-st.nygenome.org/. 26 sections were used in downstream analysis. Individual ST spots were assigned to specific spinal cord regions based on expert annotation from the original publication. Analysis of microglial signature genes was restricted to the ventral horn and dorsal horn, as the ventral horn is known to demonstrate higher motor neuron atrophy in ALS (REF). To compare levels of expression of genes between the two regions, predicted counts for all genes were normalized on a per-section basis. Next, for each gene, the summed score for all spots in the dorsal horn was compared to the summed score of all spots in the ventral horn for the same gene in the same section. Testing was conducted with Welch’s t-test with the Holm-Bonferroni correction^16,17^, setting a threshold of 0.05 for significance.

#### Identifying regulators of different microglial subtypes with CHEA3

To identify regulators, we leveraged the recently published CHEA3^18^ tool using the web interface. We performed two primary analyses: comparison of different clusters, and comparison of ALS and non-ALS microglia within individual clusters. Gene sets for input were identified as above. For the ALS vs. non-ALS analysis, we used the top 50 genes per cluster that were upregulated in ALS. Conversely, for the cross-cluster analysis, we used curated gene lists that composed of signature genes for individual clusters. All analyses were conducted using only microglial clusters. Results of the analysis per cluster were sorted by “Score”. For visualization, the top 15 predicted regulators per cluster were chosen. After removing duplicates, the Z-scored regulator scores per cluster were described in heatmaps. Clustering of regulators and clusters was done with hierarchical clustering using absolute linkage.

#### Annotation of non-microglial immune subtypes in ALS single-cell RNA-sequencing data

Non-microglial immune subtypes (clusters 10, 11, and 12) were separated from other data and re-run through a clustering/dimensionality reduction pipeline. In brief, we used SCTransform with default parameter and harmony to integrate across batches using a thetaval of 2, and used 20 principal components for downstream analysis. To identify the optimal number of clusters for downstream analysis, we used the recently described ChooseR (REF) tool to identify an optimal clustering resolution for our dataset. Next, enrichment of different immune subtypes in ALS vs. non-ALS samples and in different regions was computed as described in **Identifying differentially represented subsets between ALS and non-ALS samples** and **Identifying differentially represented subsets between regions in ALS samples**. Next, cluster-defining gene subsets were identified using the approach described in **Identification of cluster-defining gene sets.** Cluster identities were assigned with the use of canonical markers.

#### Statistical analysis and data visualization

Statistical analysis was conducted as described in the associated methods sections above. Specific p-values (both significant and not), if not found in the figures, may be found in Supplementary Information tables before and after testing for multiple correction. T-values and degrees of freedom are also provided where relevant. Unless otherwise noted, all measurements are taken from distinct samples. In general, statistical methods were not used to re-calculate or predetermine sample sizes. All plots were created in R v4.1.0 using either base R visualization packages, ggplot2^19^ with ggrepel^20^, ggfortify^21^, patchwork^22^, cowplot^23^, and ggsci^24^, or packages mentioned in the methods text. Heatmaps were made with the pheatmap^25^ package. Volcano plots were made with the EnhancedVolcano^26^ package. Figures were created using BioRender.com. All boxplots denote the 25^th^ percentile, median, and 75^th^ percentile, with whiskers representing 1.5 times the IQR in both directions. Outliers, if any, are represented as circles beyond the whiskers.

#### Data Availability

Raw scRNA-seq data (fastq files) generated from CD45+ cells isolated from ALS autopsy samples were deposited to GEO (https://www.ncbi.nlm.nih.gov/geo/) under accession number GSE204702.

#### Code Availability

Code used to perform preprocessing, clustering, cluster validation, and label transfer of scRNA-seq data in the current study are available publicly at https://github.com/jtuddenham/ALS_sc_microglia.

