## Supplemental figures and legends for "Single-cell transcriptomic landscape of the neuroimmune compartment in amyotrophic lateral sclerosis brain and spinal cord"

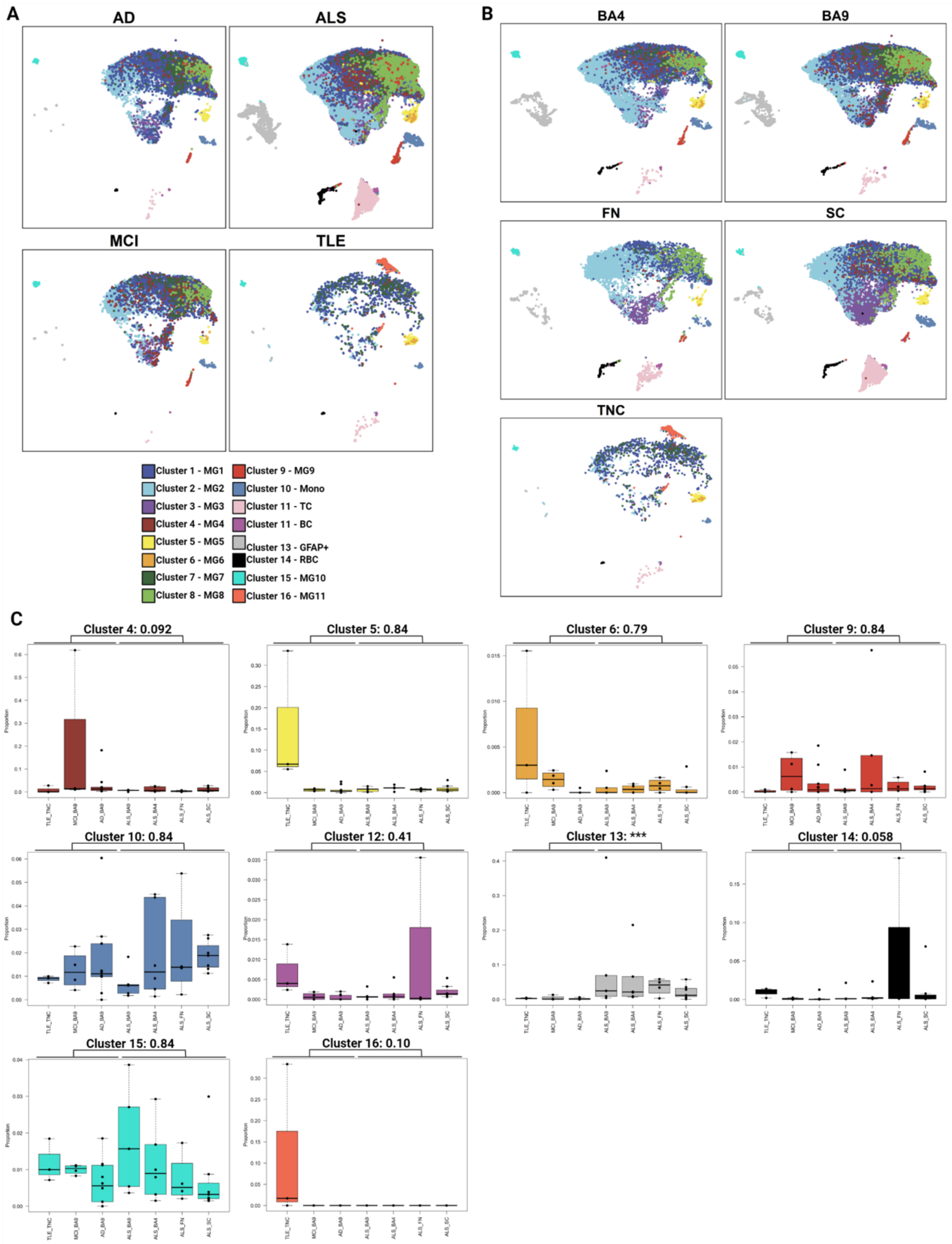


**Figure S1. Representation of human microglial subsets across diseases and regions**. Related to Figures 1 and 2. (**A**) Microglial population structure in different diseases. UMAP plots are split by disease. Overarching population structure is detected across different diseases, albeit at substantially varying frequencies. (**B**) Microglial population structure in different regions. As in (**A**), except that UMAP plots are split by region. (**C**) Remaining comparisons of subtype abundance between ALS and non-ALS samples. Boxplots as described in Figure 2B for all subtypes not shown in the prior panel. Significance testing as described previously. UMAP uniform manifold approximation and projection, AD Alzheimer's disease, MCI mild cognitive impairment, TLE temporal lobe epilepsy, ALS amyotrophic lateral sclerosis, BA Brodmann area, TNC temporal neocortex, SC spinal cord, SN substantia nigra, FN facial nucleus.

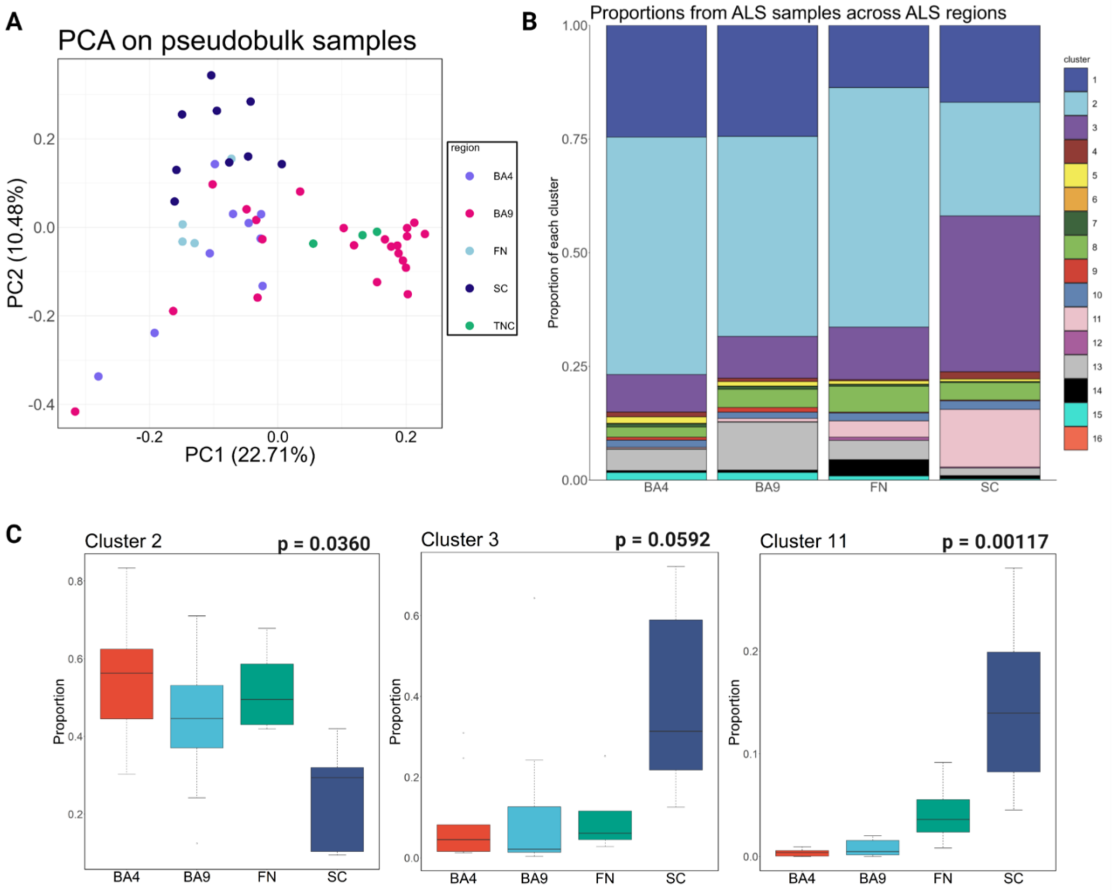


**Figure S2. Cell types and subtypes show distinct regional associations. (A) PCA demonstrates separation of pseudo-bulked samples from different regions.** PCA plot was calculated from pseudo-bulked scRNA-seq samples. Points are colored by region of origin. Notably, region represents a major component of the axis of separation between different samples, with spinal cord showing the most separation. **(B) Distributions of clusters within ALS samples.** Only ALS samples are included in this plot, and each bar shows the proportion of each subset in each of the four regions we sampled in ALS. **(C) In the spinal cord, microglial subtypes and T cells have altered abundance.** While in the ALS vs non-ALS comparison, cluster 2 is the most significantly upregulated cluster, it is significantly different between regions in ALS. In comparison, cluster 3 shows a trend towards being differentially abundant between regions in ALS, suggesting a reciprocal shift in microglial subtypes in ALS SC. Cluster 11, representing T cells, is also significantly differentially abundant between regions. Boxplots denote the 25$^{th}$ percentile, median, and 75$^{th}$ percentile, with whiskers capturing 1.5 IQR in both directions. Testing was performed with Kruskal-Wallis tests and BH correction.

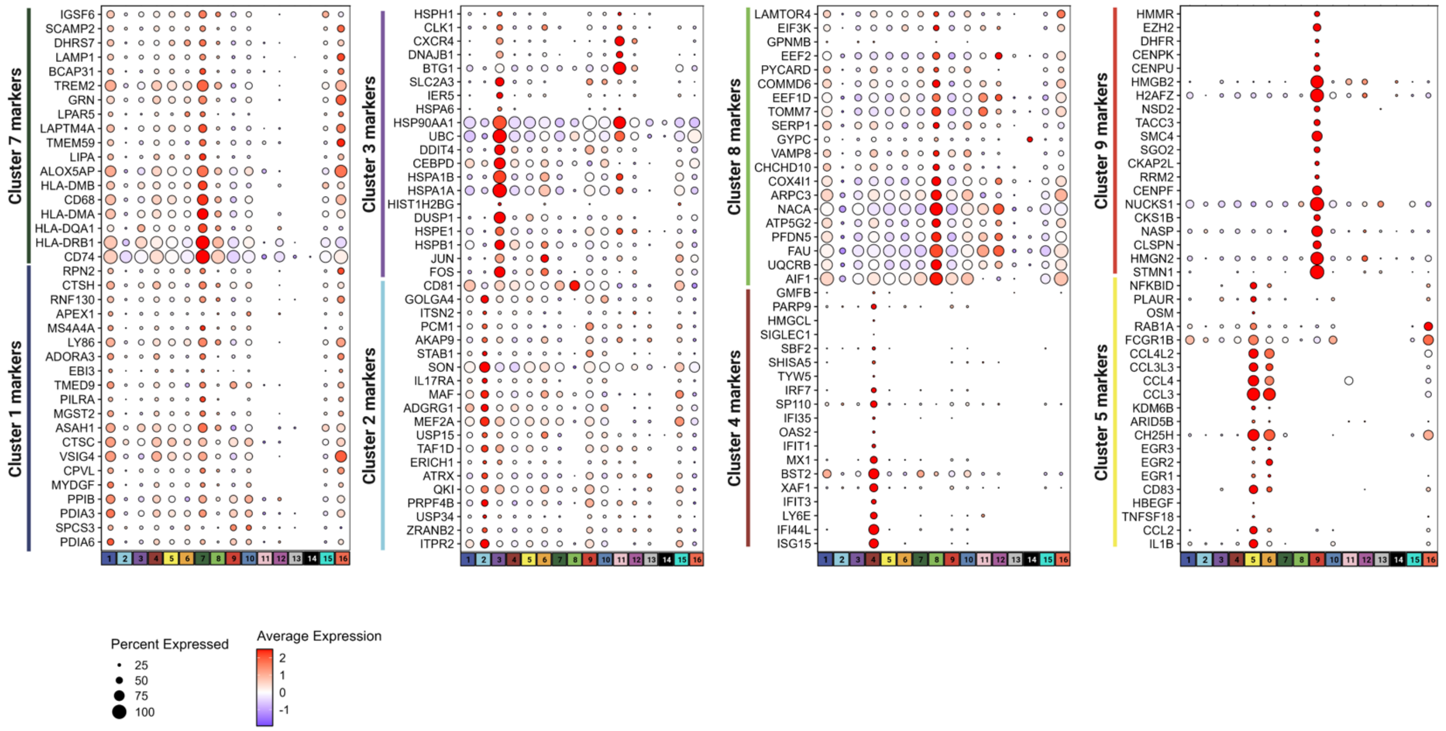


**Figure S3. Microglia subset specific gene sets.** Dot plots representing the average expression of genes (rows) identified to be enriched in microglia clusters 1 to 9 (columns). The size of the dots corresponds to the percentage of cells in which the expression of the given gene was detected in the specified cluster.

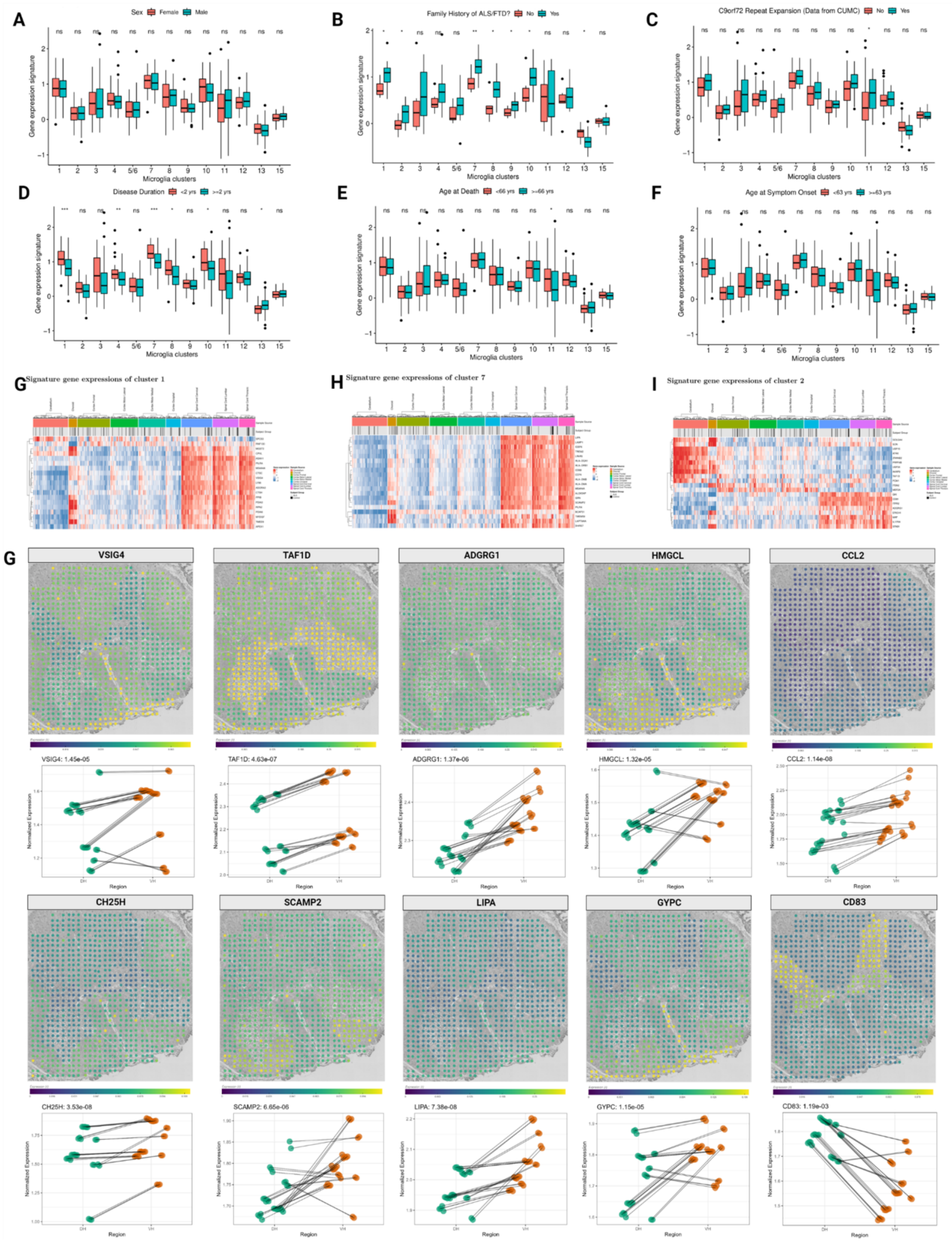


**Figure S4. Further investigation of orthogonal datasets (A)-(F) Association of cluster signatures with metadata parameters in ALS bulk RNA-seq cohort.** Dataset and analyses as in Figure 2C-D. Cluster signature correlation was analyzed in relation to: gender, ALS/FTD family history, C9Orf72 repeat expansion presence, disease duration, age at death, and age at symptom onset in the cervical spinal cord bulk RNA seq dataset. Notably, no gender differences are observed. C9Orf72 repeat presence and younger age at death both correlate with higher numbers of T cells. **(G)-(I) Regional distribution of signature genes for different clusters.** Expression of selected genes (rows) are shown across different samples (columns), separated by region. ALS or non-ALS diagnosis is denoted by the intermediate bar. Notably, clusters 1 and 7 have heavy enrichment of genes in the spinal cord, while cluster 2 signature genes show separation between spinal cord and cerebellum. (**J**) **Expression patterns of other marker genes in spatial transcriptomic data.** Plots are as described in Figure 2E/F. Notably, most marker genes are anticorrelated with MAP2 expression, but interestingly CD83 is positively correlated with MAP2 expression. TAF1D and ADGRG1 are cluster 2 markers.

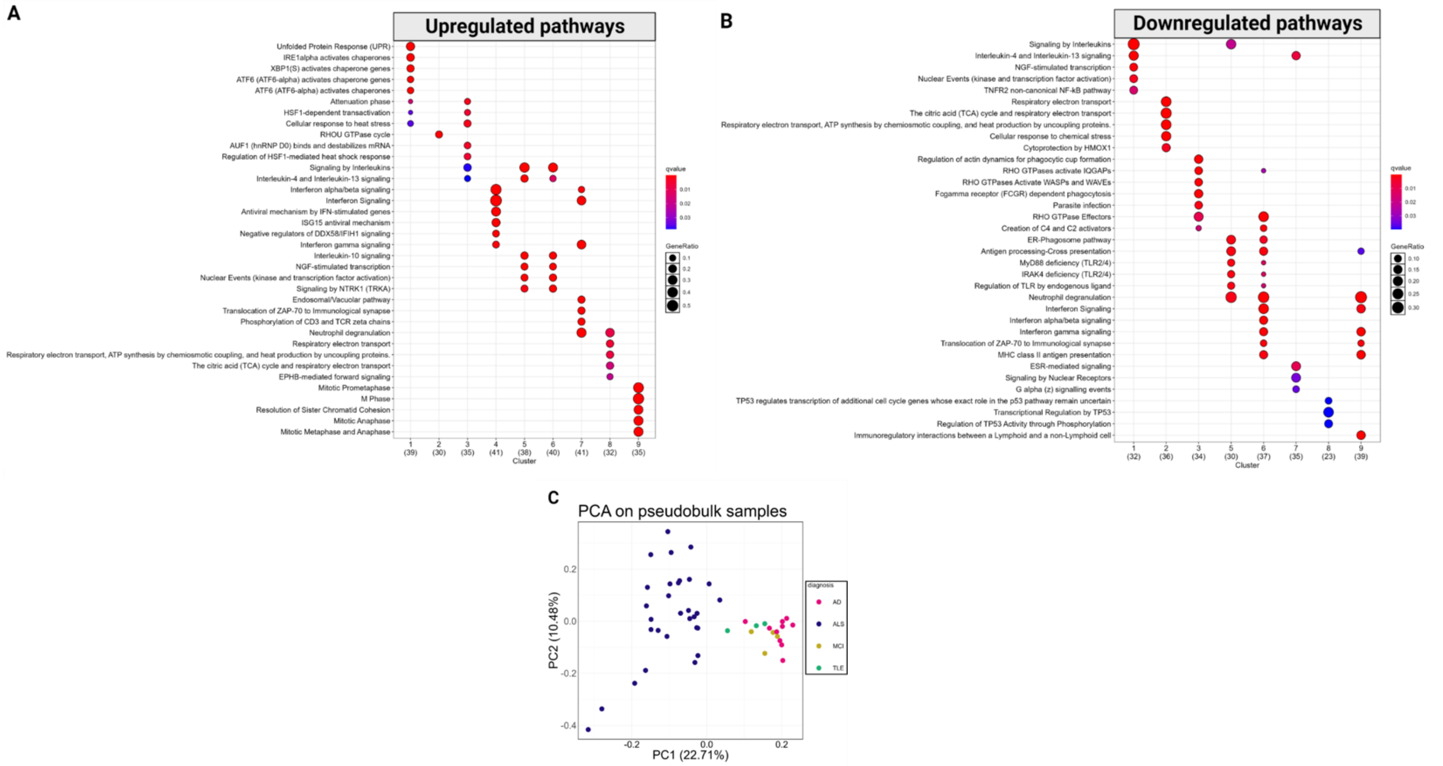


**Figure S5. Examining separation of ALS and non-ALS microglia across different data modalities.** (**A**)-(**B**) Cross-subset annotation of processes up- or down-regulated in different microglial clusters. Pathway analysis conducted with Reactome for upregulated (**A**) and downregulated (**B**) gene sets associated with each overall microglial cluster. Results displayed in the form of a dotplot, where the color is the qvalue and the size is the GeneRatio defined by the ClusterProfiler package. Results highlight differences in interleukin signaling, antigen presentation, and oxidative phosphorylation that define the different microglial subsets that are differentially represented in ALS. (**B**) ALS samples segregate from non-ALS samples in scRNA-seq. PCA plot was calculated from pseudo-bulked scRNA-seq samples. Points are colored by diagnosis. (**C**) ALS samples separate from non-ALS samples in proteomic data. PCA plot was calculated from bulk proteomic data. (**D**) ALS samples separate from non-ALS samples in ATAC-seq data. PCA plot was calculated from bulk ATAC-seq samples. Points are colored by diagnosis. (**E**) Volcano plot of ATAC-seq data.

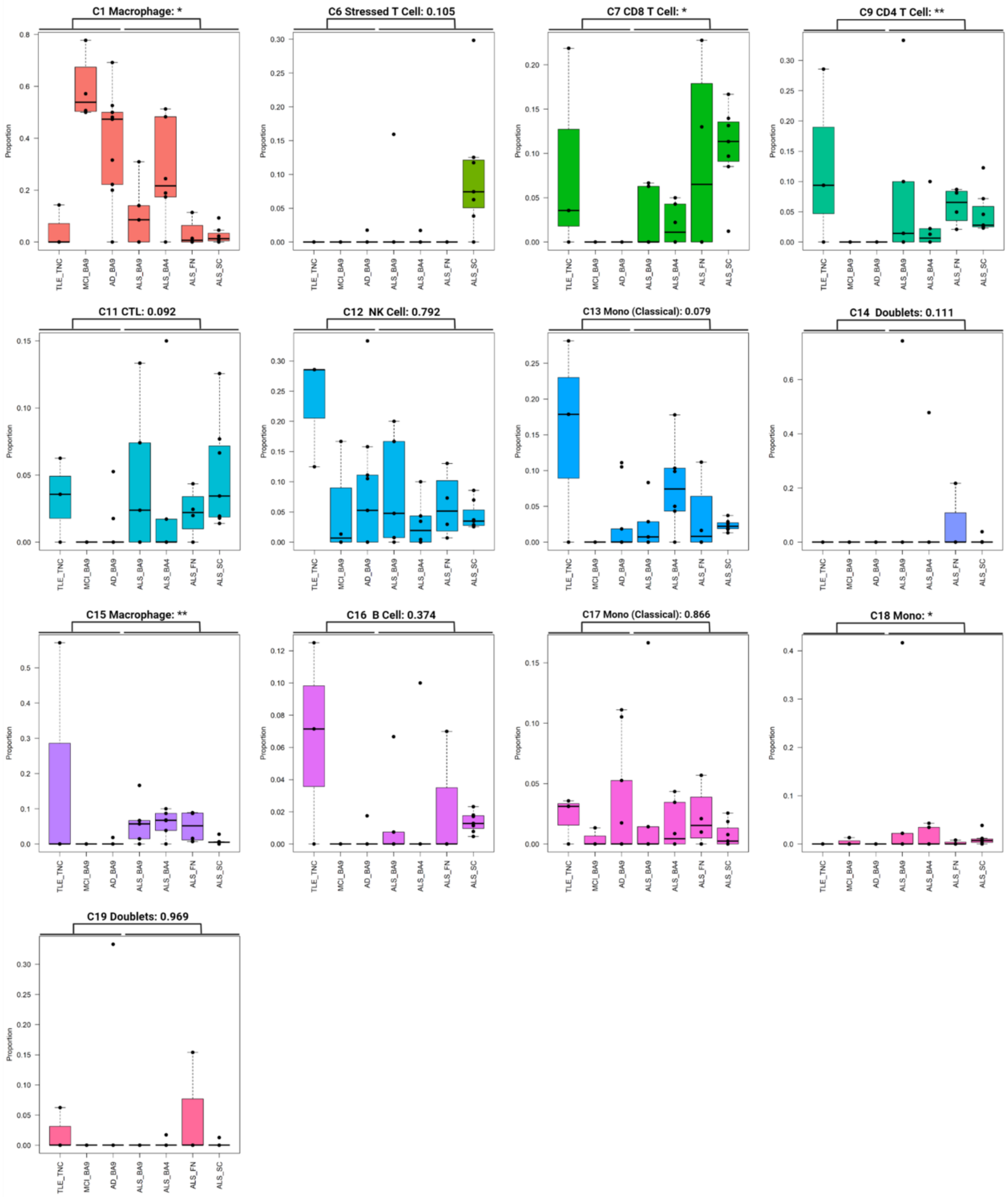


**Figure S6. Visualizing distribution of remaining immune subtypes between ALS and non-ALS samples.** Boxplots denote the 25th percentile, median, and 75th percentile, with whiskers capturing 1.5 IQR in both directions.

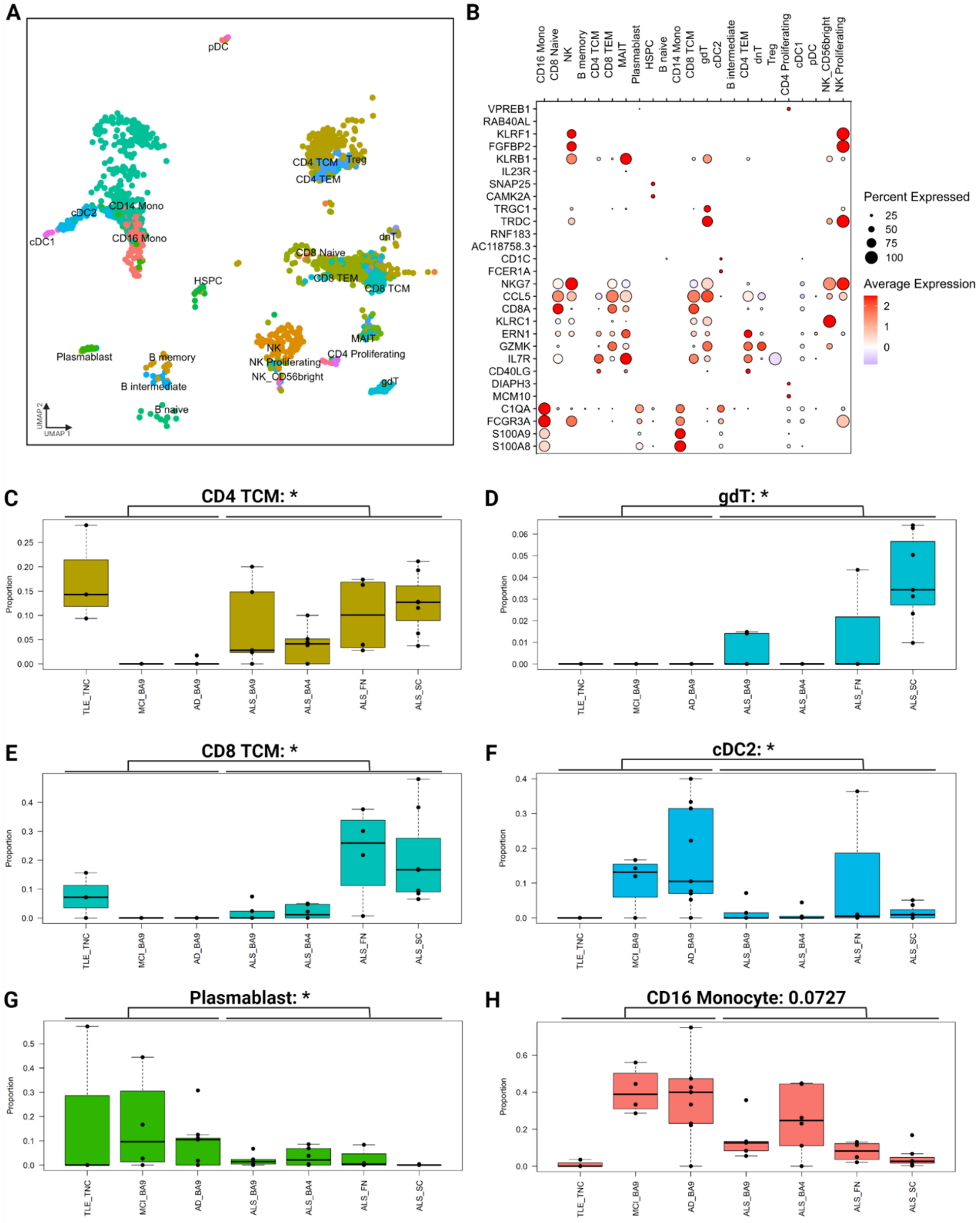


**Figure S7. Examination of non-microglial immune cells in the central nervous system by annotation with the Azimuth reference atlas.** (**A**) Annotation of non-microglial immune cells via the Azimuth PBMC reference atlas. Non-microglial immune cells were isolated in silico and separately processed. The Azimuth web interface was used to assign annotations to this dataset using the PBMC RNA/ADT reference. Resulting annotations are plotted in a UMAP plot. (**B**) Selected marker genes validate efficacy of Azimuth classification. Each column represents one of the assigned labels from the Azimuth workflow, and each row represents the z-scored expression of a given gene. The plot is colored according to level of expression and the size of each circle represents the percentage of cells in each cluster that express the gene. (**C**)-(**H**) Annotation identifies non-microglial immune populations that are differentially represented in ALS. Boxplots denote the 25th percentile, median, and 75th percentile, with whiskers capturing 1.5 IQR in both directions. Asterisks denote significance of difference between conditions, using the Wilcoxon rank-sum test and BH correction to determine significance. CD4 TCM, gDT, and CD8 TCM show enrichment in ALS, while cDC2, plasmablasts show significant depletion in

ALS. * < 0.05. ALS amyotrophic lateral sclerosis, TCM T Central Memory, gDT gamma-delta T, cDC conventional dendritic cell, NK natural killer, MAIT mucosal associated invariant T dnT double negative T HSPC hematopoietic stem and progenitor cell, pDC plasmacytoid dendritic cell.

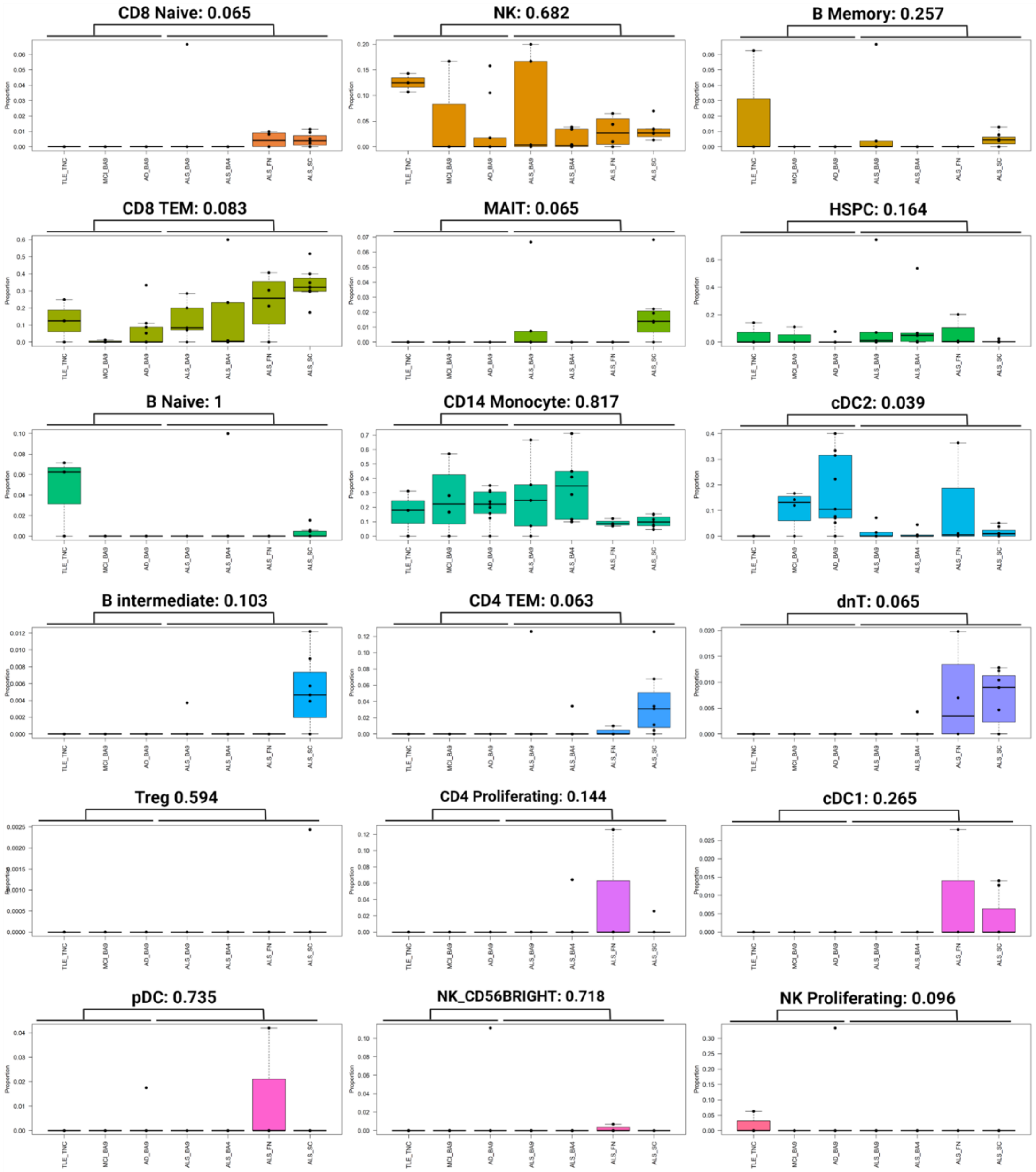


**Figure S7. Visualizing distribution of remaining immune subtypes between ALS and non-ALS samples using Azimuth annotations.** Boxplots denote the 25th percentile, median, and 75th percentile, with whiskers capturing 1.5 IQR in both directions.

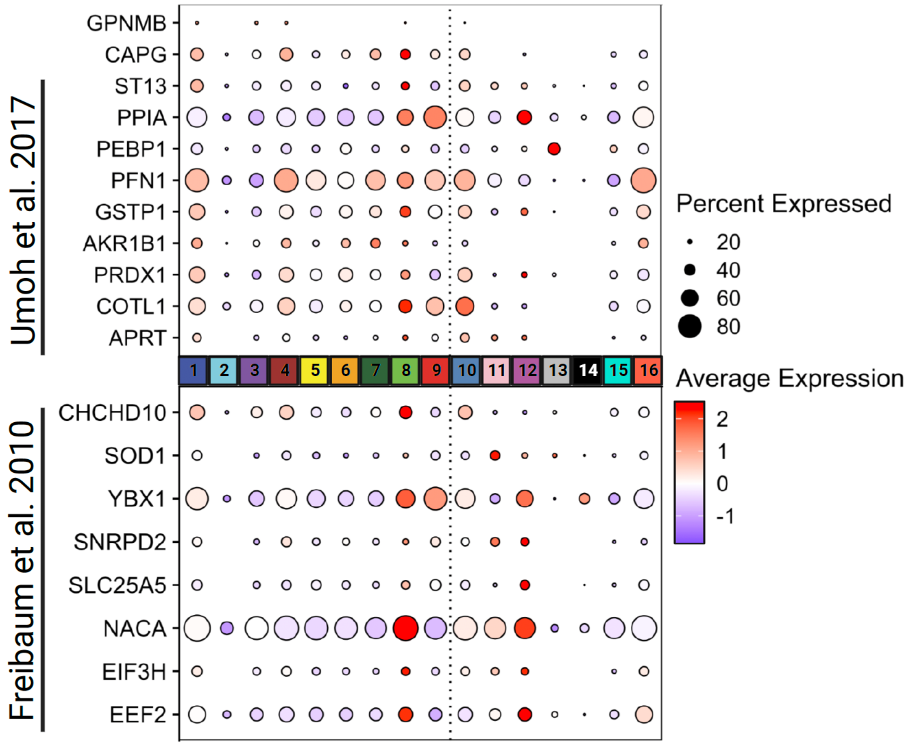


**Figure S8. Expression of ALS enriched protein module markers and TDP-43 interactors in microglia and non-microglial immune cells in ALS.** Dot plots representing average expression of genes specified on the right axis (rows) in the different clusters (columns). The size of the dots corresponds to the percentage of cells in which the expression of the given gene was detected in the specified cluster. The upper panel represents the markers from the ALS enriched protein co-expression modules from the study by Umoh et al., 2017. The lower panel demonstrates the expression of the TDP-43 interacting partners identified in Freibaum et al., 2010.

**List of supplemental tables**

**Table S1** - Overview of demographics, hashing strategy, and sequencing quality controls

**Table S2** - Pairwise marker genes across microglia clusters, related to Figure 1

**Table S3** – Cluster-vise differential representation testing between ALS and non-ALS samples and between central nervous system regions, related to Figure 2, Figure S1 and Figure S2

**Table S4** - Curated marker gene sets for individual microglia clusters, related to Figure S3

**Table S5** - Microglia subset marker gene expression changes in the dorsal and ventral horn of the human spinal cord spatial transcriptomic data, related to Figure 2 and Figure S4

**Table S6** - Cluster-vise REACTOME pathway enrichment analysis, related to Figure 1 and Figure S5

**Table S7** - Cluster-vise REACTOME analysis of up and down-regulated pathways in ALS versus non-ALS samples, related to Figure 3

**Table S8** - Cross-cluster CHEA3 analysis, related to Figure 3

**Table S9** - Cluster-vise CHEA3 analysis of ALS versus non-ALS samples, related to Figure 3

**Table S10** - De novo clustering of non-microglial immune cells

**Table S11** - Pairwise marker genes across de-novo clustered non-microglial immune cells, related to Figure 4

**Table S12** - Differential representation testing of de-novo clustered non-microglial immune cells, related to Figure 4 and Figure S6

**Table S13** - Azimuth mapping of non-microglial immune cells, related to Figure S7
